# CAnDI: a new tool to investigate conflict in homologous gene trees and explain convergent trait evolution

**DOI:** 10.1101/2023.11.18.567661

**Authors:** Holly M. Robertson, Joseph F. Walker, Edwige Moyroud

## Abstract

Phenotypic convergence is found across the tree of life, and morphological similarities in distantly related species are often presumed to have evolved independently. However, clarifying the origins of traits has recently highlighted the complex nature of evolution, as apparent convergent features often share similar genetic foundations. Hence, the tree topology of genes that underlie such traits frequently conflicts with the overall history of species relationships. This conflict creates both a challenge for systematists and an exciting opportunity to investigate the rich, complex network of information that connects molecular trajectories with trait evolution. Here we present a novel conflict identification program named CAnDI (Conflict And Duplication Identifier), which enables the analysis of conflict in homologous gene trees rather than inferred orthologs. We demonstrate that the analysis of conflicts in homologous trees using CAnDI yields more comparisons than in ortholog trees in six datasets from across the eukaryotic tree of life. Using the carnivorous trap of Caryophyllales, a charismatic group of flowering plants, as a case study we demonstrate that analysing conflict on entire homolog trees can aid in inferring the genetic basis of trait evolution: by dissecting all gene relationships within homolog trees, we find genomic evidence that the molecular basis of the pleisiomorphic mucilaginous sticky trap was likely present in the ancestor of all carnivorous Caryophyllales. We also show that many genes whose evolutionary trajectories group species with similar trap devices code for proteins contributing to plant carnivory and identify a *LATERAL ORGAN BOUNDARY DOMAIN* transcription factor as a possible candidate for regulating sticky trap development.

## Introduction

High-throughput sequencing technologies have allowed large multi-gene phylogenetic data (phylogenomic datasets) to emerge as the standard for inferring species relationships (Delsuc et al. 2005; Jeffroy et al. 2006). These datasets have become increasingly common and provide a wealth of information for investigating contentious regions in the tree of life (Dunn et al. 2008; Chiari et al. 2012; Wickett et al. 2014). Efforts to analyse the complete set of gene phylogenies in a genome, the “phylome” (Sicheritz-Pontén and Andersson 2001; Huerta-Cepas et al. 2007), are emerging as a new field.

In addition to species tree inference, these datasets also provide unprecedented opportunities to explore evolutionary history beyond species relationships. By comparing gene evolutionary histories to that of the species, it is possible to uncover patterns that reflect historical events resulting from ancient population structures (Parins-Fukuchi et al. 2021; Stull et al. 2021). In particular, these phylogenomic datasets may shed some light on the question of how similar adaptations emerge repeatedly during evolution.

Convergent evolution can be described as the evolution of phenotypic similarity (Doolittle 1994; Stern 2013; Arbuckle et al. 2014) but it can also be defined at a molecular level by considering the genetic basis of alike traits. Phenotypic convergence can reflect molecular convergence when parallel changes occur in genes underlying a given feature, but it can also arise from the recruitment of existing ancestral allelic polymorphisms or the introduction of an allele by hybridisation (either ancient or recent) (Stern 2013). Both recruitment and hybridisation are reflected in the phylome as gene tree conflict, and investigating the conflicting evolutionary histories in genomic data can provide valuable biological insights (Doyle 1992; Maddison 1997; Rokas et al. 2003). Examples of this include ancestral populations of butterflies that underwent introgressive hybridization to produce shared mimicry genes (Dasmahapatra et al. 2012), incomplete or random lineage sorting resulting in unusually high levels of homoplasy in cacti (Copetti et al. 2017), the distribution of karyotypic features in mammals (Robinson et al. 2008), and the horizontal transfer of large DNA regions that explains the phylogenetic distribution of C4 photosynthesis in grasses (Dunning et al. 2019). An adaptive role for genes in conflict was also established in tomato plants (Pease et al. 2016), and more recently in Marsupials (Feng et al. 2022) and Diatoms (Roberts et al. 2023). Hence, investigating conflicting patterns can illuminate the evolution of complex traits and uncover molecular players that could contribute to novelty.

To capture the true extent of genome-wide patterns of discordance and all relevant biological signals, it is essential to analyse as much of the phylome as possible; and currently, the most cost-effective approach to accessing nearly complete phylomes is through transcriptomes. However, current studies and methods for investigating conflict often focus on analysing orthologs inferred from homolog trees. Focusing solely on the inferred orthologs results in the loss of information and potentially interesting biological signals. Paralogs (gene copies that arise within a population due to gene duplication) must still, however, be accounted for, as they can mislead both inferred species trees and gene tree conflict analyses.

### A case study in phenotypic convergence: the carnivorous Caryophyllales

The flowering plant order Caryophyllales contains several emblematic species of carnivorous plants, including the Southeast Asian pitcher plants, sundews, and the Venus flytrap. While carnivory is the predicted ancestral condition in the clade of all carnivorous Caryophyllales (Cameron et al. 2002; Heubl et al. 2006; Renner and Specht 2011), not all species in this clade are carnivorous. The Ancistrocladaceae family, for example, is entirely non-carnivorous.

The clade of carnivorous Caryophyllales exhibits several instances of apparent phenotypic convergence. Trap type and stem morphology are two particularly interesting examples of major, complex adaptations involving many genes. Underlying both is the placement of the clade of Ancistrocladaceae and Dioncophyllaceae as sister to the monotypic family Drosophyllaceae, a relationship supported by phylogenomic datasets composed of hundreds or even thousands of genes (Walker et al. 2017, 2018; Baker et al. 2022). This inferred relationship is of note as the families Ancistrocladaceae and Drosophyllaceae exhibit striking morphological similarities not with each other but with other, more distantly related, species in the clade.

Specifically, Drosophyllaceae, a family containing only one species, the Portuguese sundew (*Drosophyllum lusitanicum*), is not sister to the genus *Drosera* (commonly known as the sundew family) even though both have a sticky trap mechanism of capturing prey and a herbaceous stem morphology. Moreover, there is also *Triphyophyllum peltatum*, a liana in the Dioncophyllaceae, which is native to West Africa and develops sundew like carnivorous sticky trap leaves under phosphorus deficient conditions (Winkelmann et al. 2023).

This polyphyletic distribution of sticky traps within the carnivorous Caryophyllales indicates that either true convergence has occurred or a precursor to the sticky trap existed ∼75 mya, with the extant phylogenetic distribution of these traps resulting from incomplete lineage sorting, horizontal gene transfer, or introgression.

Another implication of Drosophyllaceae being sister to the clade of Ancistrocladaceae and Dioncophyllaceae is that this relationship places the lianas of Ancistrocladaceae as more closely related to the herbaceous Drosophyllaceae rather than the carnivorous pitcher plant family Nepenthaceae, which is also a liana. This relationship would indicate that either an ancestral population that existed approximately 67 mya exhibited a woody stem morphology later lost for Drosophyllaceae or that such a lifestyle has been gained twice independently (Cameron et al. 2002; Heubl et al. 2006; Walker et al. 2017).

Here we introduce CAnDI (Conflict And Duplication Identifier), a new program we developed to analyse all gene relationships in an unbiased way and investigate conflict across the phylome. This program dissects all relationships within homolog trees and analyses them for conflict, concordance, or origination by duplication. This method expands upon previous work in conflict detection, as it analyses and reports every instance of conflict in a dataset and allows users to search for conflicts of interest. In addition, it allows users to search for patterns of interest within homolog trees. We demonstrate that CAnDI can identify evolutionary patterns that cannot be detected using current approaches by applying it to a series of datasets from various closely and distantly related groups. We further show that the use of orthologs accurately demonstrates the amount of conflict expected from the whole phylome. Following this, we conduct a more detailed case study in the carnivorous Caryophyllales, where our new method enables us to provide genomic evidence that molecular variants shared by extant species with a mucilaginous sticky trap were already present in the ancestor of all carnivorous Caryophyllales. Using CAnDI, we also identify a set of genes that could have participated in the emergence of the sticky trap morphology during the evolution of the Caryophyllales. In particular, our analyses, combined with quantification of gene expression levels in sundew tissues, single out the transcription factor *LBD4-like* as a candidate regulator of sticky trap development.

## Materials and Methods

### Comparison of whole homolog trees with ortholog trees for identifying gene tree conflict in six datasets

We compiled six datasets of species trees, ortholog trees, and gene families (homolog trees) from published studies that broadly span the tree of life. We used the species relationships and homolog trees inferred in the initial studies as a basis for each conflict analysis. If available, we used rooted homolog trees from the original study. If the rooted homolog trees were not available, we used the extract_clades.py script from (Yang et al. 2015) to extract them.

All of the datasets, both homologs and orthologs, were analysed using CAnDI to count the total number of nodes assessed corresponding to each node in the species, where each node was assessed as either conflicting with the species tree or concordant with the species tree. Gene duplication nodes were not counted.

We extracted 12,172 rooted homologs from the unrooted homologs previously inferred for the plant family Amaranthaceae (AMAR) by (Morales-Briones et al. 2021), using a threshold of 40 ingroup taxa. The AMAR dataset did not have support values for homolog trees. The 12,699 orthologs of the AMAR dataset with appropriate outgroup representation for rooting were from the original study where they were extracted using the monophyletic outgroup (MO) method (Yang and Smith, 2014). The Diatom (DIAT) dataset from (Parks et al. 2018) had duplications extracted with a minimum of 10 ingroup taxa, resulting in 18,914 homolog trees. The support values for this dataset were from a rapid bootstrap analysis; thus, we applied a cutoff of ≥70%. The 197 orthologs from the original study, that were extracted using the Rooted Tree (RT) method (Yang and Smith, 2014), were used for the ortholog comparison for the DIAT dataset. The Legumes dataset (LEGU), consisting of 8,038 homolog trees from (Koenen et al. 2020), had been extracted in the original study, and support values on the nodes were from a rapid bootstrap analysis; thus, a cutoff of (≥70) was used. For the LEGU orthologs, we used the ortholog trees extracted with the RT method from the original study. To extract the homologs we used the same outgroup taxa, resulting in 1014 trees with outgroup representation. The Hymenoptera dataset (HYMN) was originally published by (Johnson et al. 2013) but was subsequently re-analysed by (Smith et al. 2015), who identified 5,863 homolog trees and calculated rapid bootstrap support; therefore, we used a support cutoff of ≥70% for the HYMN dataset. To generate orthologs for the HYMN dataset, we used the RT method with a minimum cutoff of four taxa using the prune_from_rooted_tree.py script of (Morales-Briones et al. 2021), resulting in 6519 orthologs. The 6902 homolog trees from the Ericales (ERIC) dataset from (Larson et al. 2020) were analysed using a support value of (SH-aLRT ≥ 80). Conflict analysis on ortholog trees from the ERIC dataset was conducted using the 382 rooted ortholog trees from the original study that had been extracted using the RT method. The details of the carnivorous Caryophyllales (CARN) dataset, also used in this comparison, are described below.

### Investigating the relationship of conflict with trait distribution in the carnivorous Caryophyllales (CARN) dataset

We used a minimum threshold of four ingroup taxa to extract 6,006 rooted homolog trees from the unrooted homolog trees inferred by Walker et al. (Walker et al. 2017). We term this the carnivorous Caryophyllales (CARN) dataset. Information regarding this and other datasets as well as all supplementary material is available on DRYAD (doi:10.5061/dryad.g4f4qrfzq). The 1237 orthologs we used for the CARN dataset were those from the original study, which were extracted using one-to-one orthology with 100% gene occupancy required. All trees in the CARN dataset had SH-aLRT support values; thus, we used a support cutoff of ≥80%. We then used CAnDI to identify all homolog trees in the CARN dataset that contained relationships (bipartition) of interest and the number of times the relationship of interest occurred in a single homolog tree. We identified trees with the bipartition *Drosophyllum* + *Drosera* and all trees with the bipartition *Drosophyllum* + *Ancistrocladus*. We also used CAnDI to identify all bipartitions that contained only *Nepenthes ampullaria*, *Nepenthes alata,* and *Drosophyllum lusitanicum*. The same procedure was performed to identify all instances in homolog trees where Ancistrocladaceae (Ancistrocladus) was sister to Nepenthaceae (*N. alata* and *N. ampullaria*). For each group of taxa investigated, we compared the number of bipartitions that contained only those taxa against the number of instances where an additional species was also included in the bipartition to eliminate any bias caused by differences in gene recovery in different samples.

### Investigating gene function in the CARN dataset

Annotations for the CARN dataset were inferred using the coding sequences of the genome of *Arabidopsis thaliana* (TAIR10; Downloaded from Ensembl Plants on Jan 27^th^, 2021). The best blast hit (e-value 1e-3) to the transcriptome of *Nepenthes alata* from each homolog cluster was used to predict the function of each cluster. Using this approach, 137 out of the 146 clusters supporting the monotypic family Drosophyllaceae sister to the genus *Drosera* could be annotated (9 clusters failed to return a blast hit against *A. thaliana*. These 137 annotations represented 132 distinct Arabidopsis genes. Gene ontology analyses with PANTHER16.0 were performed on this list of 132 *A. thaliana* genes using the PANTHER Overrepresentation Test (Released 20210224) and the GO Ontology database (DOI: 10.5281/zenodo.5228828 Released 2021-08-18) and either the “GO biological process complete” or the “GO molecular function complete” dataset. The Fisher’s exact test was performed with False Discovery Rate correction for each analysis. A *Chitinase-like protein 1* [homolog to *Arabidopsis CTL1*; labelled cluster2518 by Walker et al. (Walker et al. 2017)] was further investigated since this gene is associated with carnivory. We spot-checked the annotation for this gene by testing the blast annotation of other homologs within cluster2518 against the NCBI RefSeq non-redundant protein database to ensure that this was not the result of a spurious hit. All sequences from cluster2518 matched a Chitinase gene. We then performed the reverse mapping feature with CAnDI (“-r”) and mapped the coalescent-based species tree topology for the CARN dataset to the cluster2518 homolog tree.

### Plant material

*Drosera admirabilis*, *Drosera aliciae* and *Drosera coccicaulis* were purchased from Wack’s Wicked plants (UK), grown in ambient conditions (natural light, with temperature kept between 18-25°C) and watered from the base with distilled water. Tissues were harvested from emerging young leaves (unfurled primordia <1cm long, trichomes absent or not yet able to secrete mucilage), mature leaves representing fully developed sticky traps densely covered with trichomes (stalked glandular hairs) secreting mucilage droplets (Fig. S5) and inflorescences and immediately frozen in liquid nitrogen. For each species tissues were harvested from three distinct individuals to constitute three biological replicates for the gene expression analysis. All tissue samples were stored at −80°C until RNA extraction.

### Gene expression analysis

Frozen tissues (from three distinct individuals for each species, constituting three biological replicates) were ground to a fine powder using a mortar and pestle. RNA was extracted using the Spectrum Plant Total RNA kit (Sigma-Aldrich) and retro-transcribed using SuperscriptIII reverse transcriptase (Invitrogen) following the manufacturer’s instructions. Quantitative real time PCR was performed using the Luna Universal qPCR Mastermix (NEBiolabs) and the Light Cycler 480 system (Roche). For each species and tissue type, three independent biological replicates were used (i.e., RNA extracted from three distinct individuals) and three technical replicates were performed for each condition (i.e., three repeats of the qRT-PCR reaction assessing gene expression for a given tissue type, of a given biological replicate for each species). *Actin* and *eIF4A* homologs were used as reference genes as established by Arai and collaborators (Arai et al. 2021). The primer efficiencies were 100% for *Actin*, 93% for *eIF4* and 109% for *LBD4-like* homolog genes. Primers were designed to match gene regions conserved between the three different species of *Drosera*. Primer sequences are given in the Supplementary Spreadsheet 3. The gene expression was calculated relative to the housekeeping gene actin and the common base method, which accounts for the measured efficiency of each primer pair, was used to calculate relative expression levels (Ganger et al. 2017). Welch’s t-tests or ANOVA followed by Tukey’s HSD post-hoc tests were conducted in R [https://www.r-project.org, (R Core Team 2017)] and used to test for the likelihood that LBD4-like average expression levels were equivalent across tissues.

## Results and Discussion

### CAnDI provides a procedure to dissect relationships and analyse conflict in whole homolog trees

Homolog trees contain all gene sequences with similarities resulting from their shared evolutionary history, such as those originating through gene duplication (paralogs) or speciation (orthologs). Gene duplication can occur due to local tandem duplications, chromosome duplications, or genome-wide duplication events (Lynch and Force 2000). Ancient whole genome duplications are particularly widespread in plants due to their tolerance for polyploidy (Clark and Donoghue 2018), meaning patterns of gene duplication and loss particularly characterize many plant gene families. However, gene duplication, by any mechanism, is also widespread in other eukaryotes, and gene family analyses are becoming more popular throughout evolutionary biology (Iñiguez and Hernández 2017; Copley 2020). Hence, homolog trees widen the available data for assessment and analysis of phylogenetic conflict compared to approaches that focus on only ortholog trees. With this and the widespread nature of gene duplications in the eukaryotes in mind, methods that detect and account for gene duplications on a gene family tree, allowing for the analysis of the whole homologous gene tree and not just a part of it, have widespread applicability.

Existing approaches for exploring conflict in large, multi-gene phylogenetic datasets include counting gene concordance frequencies (Ane et al. 2006; Baum 2007; Smith et al. 2015; Minh et al. 2020), dividing the tree into quartets (Pease et al. 2018; Zhou et al. 2020), and comparing gene tree likelihoods either through Bayes factors or gene-wise likelihoods (Gatesy and Baker 2005; Brown and Thomson 2016; Shen et al. 2017; Walker et al. 2018). However, these approaches tend to rely on inferred orthology. This is due to the fact that any homolog tree containing a duplication could not be analysed in entirety for conflict or concordance with a species tree, with the existence of multiple tips from the same taxon confounding this process. Previous analyses have used various orthology inference methods to mitigate such a loss of information (Zmasek and Eddy 2002; Hejnol et al. 2009; Yang and Smith 2014). However, all previous methods discard at least some information about speciation events that occur post-duplication, meaning many speciation nodes are not assessed for conflict even though they may be informative.

To perform conflict analyses on entire homolog trees, we developed the novel program CAnDI (Conflict And Duplication Identifier, https://github.com/HollyMaeRobertson/CAnDI). This program allows for the analysis of conflict in multiple ways. Firstly, as in PhyParts, (Smith et al., 2015) rooted homolog trees and ortholog trees can be mapped to a rooted species tree, generating count data for conflicts and concordances at each node in the species tree. In addition to this, a rooted species tree can also be mapped to a rooted homolog tree, labelling each node in that homolog tree as one of three mutually exclusive categories: a concordant node, a conflicting node, or a duplication node. In either case, analysis of whole homolog trees requires the specification of each node in each tree as one of either a duplication or a speciation node. Duplication nodes are those where sister clades contain at least one overlapping taxon, resulting from gene duplication. All non-duplication nodes, inferred to have arisen by speciation, can be analysed for conflict or concordance with the species tree.

In order to conduct such an analysis, CAnDI decomposes the homolog tree into subtrees by the following process (Fig. 1): first, every child node of a duplication node is copied into a list of subtrees along with its downstream structure (Fig. 1A-B). Then, in any subtree itself containing a duplication, that duplication node is collapsed (Fig. 1C): every downstream node is removed, and the node is marked with a non-repeating list of all the taxa originally present. This ensures that each speciation node is only analysed once: otherwise, a speciation node downstream of (for example) two duplications would be analysed twice, once in the larger subtree originating at the first duplication and a second time in the subtree originating at the second duplication. Following the collapsing of duplications, any subtrees where the root node leads directly to a tip (i.e., the tree contains one taxon) and any subtrees where the root node is a duplication (all speciation nodes in these trees will be analysed in the two child subtrees of that duplication) are excluded (Fig. 1D), leaving a set of subtrees that form a nonoverlapping, complete description of every speciation node in the original homolog tree in a form that can be analysed using a bipartition method (Fig. 1E) (Salichos and Rokas 2013).

**Figure 1.**
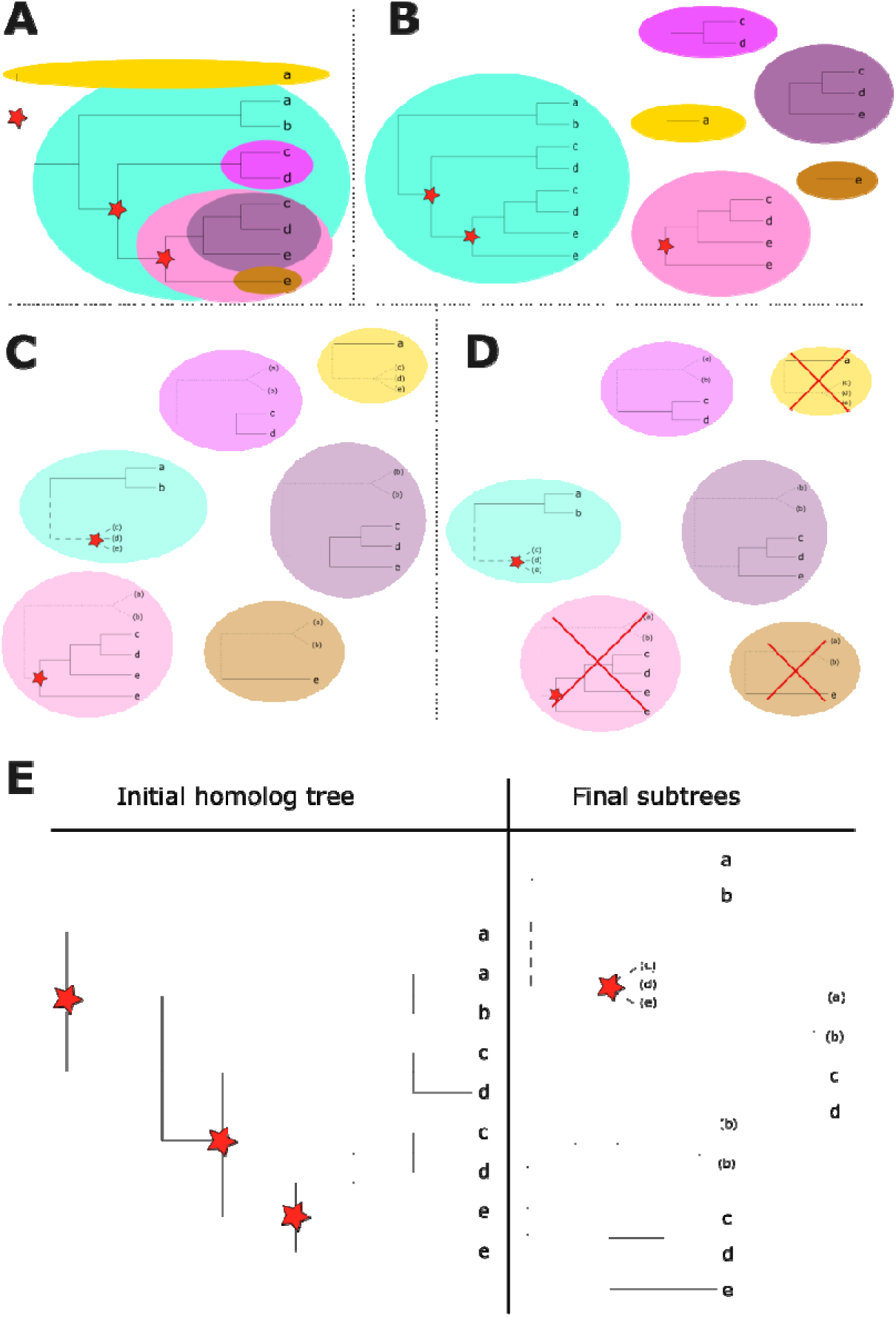
Overview of the decomposition process of a whole homolog tree into subtrees. **A-D**: Steps required for subtree decomposition and subsequent processing. The homolog tree, containing homologs from 5 distinct species (a, b, c, d and e), is first decomposed into preliminary subtrees by splitting at each duplication node (red stars) (**A**), with every child node of every duplication in the tree being added to the subtree list (**B**). Any duplications located within each of these subtrees (except at the root node) are collapsed and marked as not to be analysed (dashed branch as in the leftmost center subtree, **C**). The outgroups of each subtree are also included but labelled as not to be analysed to enable searching for missing taxa (dotted branches, **C**). Any subtrees whose root node is a duplication or leads directly to a tip are finally excluded (**D**), leaving only the informative subtrees with no overlap. **E**: The initial homolog tree and the final subtrees to be analysed are shown side-by-side.

Bipartitions form the basis for conflict and concordance in the same manner as PhyParts (Smith et al. 2015). As relationships arise from both speciation and duplication in the homolog tree, the left side of the bipartition (ingroup subtree) is evaluated against all bipartitions in the species tree (Fig. 1). The right side of the bipartition from the homolog tree is all taxa found before the duplication and any taxa within the ingroup subtree that are not in the left side of the bipartition. Any taxa in the species tree not found in either the left or right side of the homolog tree bipartition are removed from the species tree bipartitions to account for missing data. Concordance is identified as a species tree bipartition that perfectly matches a homolog tree bipartition. Conflict occurs when a homolog bipartition overlaps with a species tree bipartition but contains a non-matching set of taxa. If a support value is specified, bipartitions in the homolog tree with support below that value are not evaluated and are treated as uninformative.

CAnDI also identifies alternative conflict: under some circumstances, two non-nested nodes in a single subtree can conflict with the same node at the species tree. A single node from the homolog tree can also conflict with multiple species tree nodes. In the former case, both conflicting nodes are reported, but only one instance of conflict is counted. In the latter case, CAnDI reports the node at which conflict first occurs. Finally, CAnDI allows users to search for a particular relationship of interest. This searching procedure first creates subtrees from each homolog tree as above, then searches for the bipartition corresponding to the conflict of interest in the subtrees. How often a given homolog tree has the conflict and how many homolog trees have the same conflict can both be reported.

Whole-homolog approaches and methods for analysing the phylome have been used for other kinds of analysis than conflict [reviewed by (Smith and Hahn 2021)]. Entire homolog trees have been shown to provide a holistic approach to species tree reconciliation without the need to pre-infer orthology and are capable of recapitulating the same species relationships as using only orthologs (Smith and Hahn 2021). ASTRAL-Pro, a coalescent-based approach to infer species relationships, uses a quartet tree decomposition strategy that separates speciation and duplication nodes (Zhang et al. 2020).

Furthermore, two approaches have been developed to decompose the homolog trees and analyse them with respect to species relationships. Both methods use bipartitions as the basis for the analysis. The TreeKO method (Marcet-Houben and Gabaldón 2011), implemented in ETE3 (Huerta-Cepas et al. 2016), calculates the Robinson-Foulds (RF) distance between a species tree and a homolog tree(s). The RF distance involves decomposing each tree into bipartitions and counting the total number of bipartitions that are unique to each tree. As this involves separating duplication from speciation nodes, the tree dissection approach is similar to the one utilized here [e.g., the entire phylome (Sicheritz-Pontén and Andersson 2001)]. The biggest difference is the TreeKO method only provides a support score for the species tree. In contrast, the gene search approach in CAnDI allows the user to identify specific relationships in gene trees. CAnDI is also designed for detailed analysis of homolog trees, such as mapping species relationships onto the homolog tree and providing the number of conflicting relationships with a given species tree node.

The summary of gene trees from CAnDI is most similar to the second method, Phyparts (Smith et al. 2015). Phyparts takes a homolog tree-centric approach that involves decomposing the entire homolog tree into its respective bipartitions. The decomposition allows the program to determine whether the homolog tree conflicts or not. However, it bypasses the orthology detection procedure entirely by stipulating that each homolog tree can only be concordant, conflicting, or both at a single time with the species tree. Furthermore, as Phyparts is not designed for searching for specific conflicts, the bipartitions focus on the deepest conflicting bipartition in the tree as a summary of conflict. In short, this means that an individual homolog tree will not be summarized as having multiple conflicts or concordances.

TreeKO and PhyParts allow users to input species trees, but only to map homolog trees onto a species tree, rather than also allowing users to map the species tree onto a homolog tree. The method we developed for this study and implemented in CAnDI enables users to investigate all conflicts and concordances from a single homolog tree, unlike previous methods that can only infer whether a tree is conflicting and not the number of nodes at which an individual homolog tree is concordant or conflicting. However, we stress that Phyparts and TreeKO function for their respective purposes, and the differences outlined are not criticisms but explanations of how CAnDI has built upon these authors’ work.

### Using homolog trees as a general basis for exploring conflict allows for more comparisons

We applied CAnDI to datasets from six distinct groups of eukaryotes (hymenopterous insects, unicellular diatoms, the carnivorous Caryophyllales and three other groups of flowering plants) to compare conflict analysis based on homolog trees to that of inferred ortholog trees. We found that using entire gene families allowed significantly more relationships to inform the analysis (Fig. 2). As the orthologs from the original study were used, if the study required 100% taxon occupancy for all genes then each node was analysed the same number of times.

**Figure 2.**
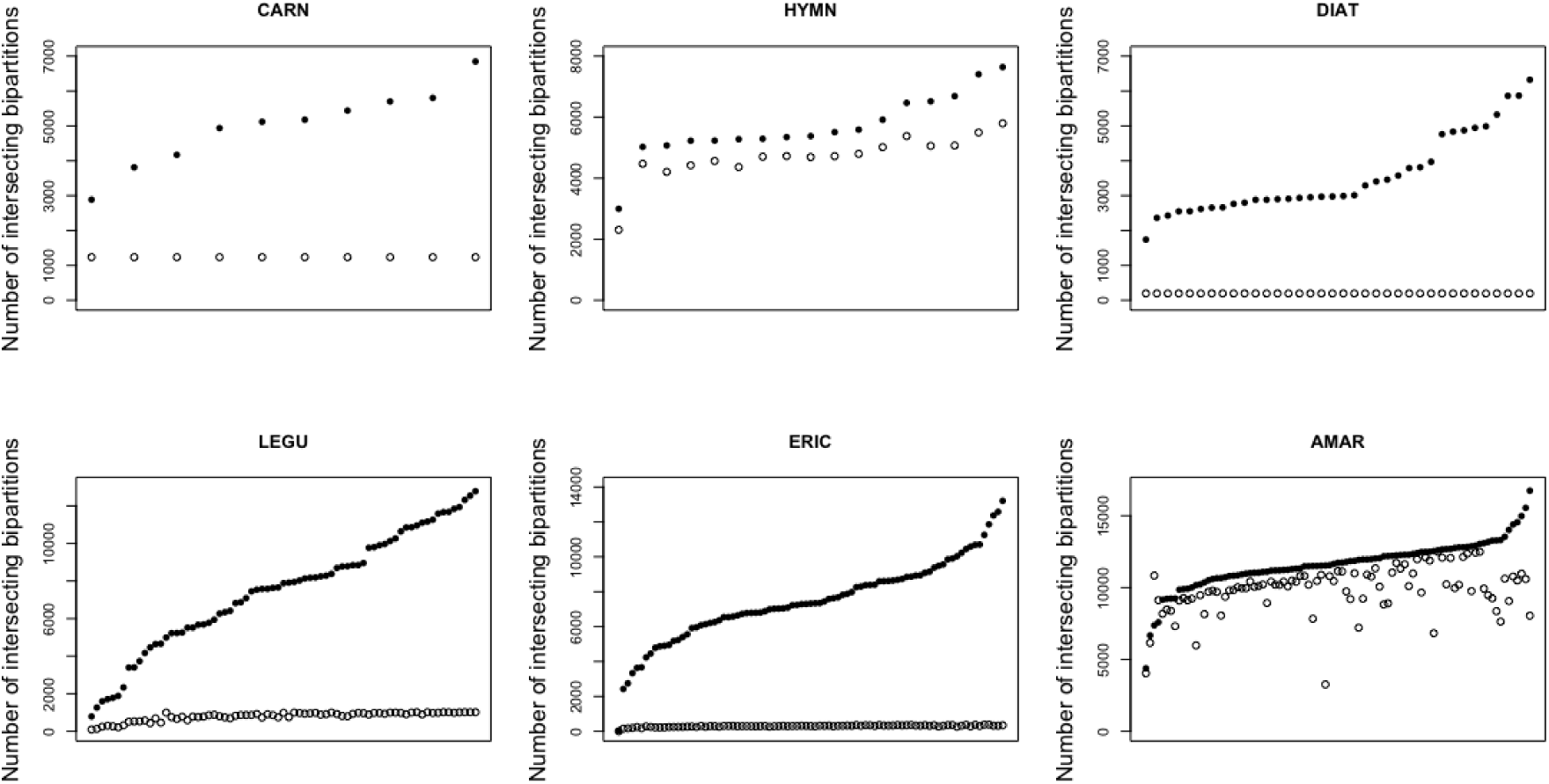
Using entire gene families provides more relationships to inform analyses and uncover conflicts. The graphs correspond to datasets from six distinct groups: the Hymenoptera order of insects (HYMN); the diatoms, a lineage of microbial eukaryotes (DIAT) and four different plants groups: the carnivorous Caryophyllales (CARN), Fabaceae (LEGU), Ericales (ERIC) and Amaranthaceae (AMAR). The graphs show the ortholog bipartitions in the species tree, ordered along the x-axis according to the number of intersecting bipartitions (fewest to most). The filled in circles are the number of times the bipartition is analysed using homologs, and the open circles are the number of times the bipartition is analysed using orthologs.

In these analyses, we extracted rooted homolog trees using the same approach as Yang and colleagues (Yang et al. 2015). This approach required a cutoff for the minimum number of ingroup taxa. However, recent work on non-time reversible models should allow users to extend even further rooting homologs without the need to extract duplicates from the homolog tree. Homolog trees rooted in this way extend the method, allowing users to analyse the entire phylome for any taxa with a sequenced genome.

Using inferred orthology is not an issue for species tree inference (Smith and Hahn 2021) or identifying processes associated with speciation patterns among genes (e.g., incomplete lineage sorting), where conflicts of interest are usually the most common ones. However, this information loss can be costly when investigating topologies that rarely arise, such as in this study, where we compare the relationship between trait distribution (distinct trap or stem morphologies) and conflict. Moreover, relying on inferred orthology is often insufficient for studies focusing on one gene family, especially in cases where that gene family has undergone significant expansion in multiple lineages. The approach we implemented in CAnDI provides a solution to overcome those limitations.

### Evolutionary conflict helps explain trait distribution in the carnivorous Caryophyllales

To further investigate the potential value of CAnDI as an analytical tool, and with the specific goal of identifying adaptive conflict, i.e. those specific conflicts which appear to correspond with a phenotype of interest’s distribution on the species tree, we turned to the carnivorous Caryophylalles.

Existing consensus (Cameron et al. 2002; Heubl et al. 2006; Renner and Specht 2011) states that the last common ancestor of the carnivorous Caryophyllales clade was carnivorous. As a precaution, we revisited this hypothesis with increased taxon sampling (Supplementary Methods) and our analysis predicted an ancestor of all the carnivorous Caryophyllales with a sticky trap (Fig.S1). This is consistent with the findings of Renner and Specht (Renner and Specht 2011), who used evidence from the different types of glands present across the carnivorous Caryophyllales to argue that the snap traps of *Dionaea muscipula* and *Aldrovanda vesiculosa* evolved from a simpler modified leaf similar to the modern sticky trap. A similar evolutionary path could also account for the emergence of *Nepenthes* pitcher traps. All carnivorous traps within the Caryophyllales are modified leaves, but snap traps and pitfall traps both exhibit elaborated morphologies, less reminiscent of a typical leaf than the sticky traps of genus *Drosera* or the species *Drosophyllum lusitanicum* (monotypic family Drosophyllaceae) and *Triphyophyllum peltatum* (family Dioncophyllaceae). However, this remains a claim made on phenotypic evidence, which can be misleading (Corbett-Detig et al. 2020), with no investigation of the evolutionary histories of the genes contributing to sticky trap development. Regardless of the mechanisms behind the subsequent elaboration of the pitfall and snap trap forms from an ancestral sticky trap, our results indicate that the presence of a sticky trap in unrelated sundews is unlikely to be a true molecular convergence (i.e., independent molecular basis) but could instead be the product of introgression or incomplete lineage sorting. Thus, the presence of the sticky trap in phylogenetically distant lineages is likely an example of hemiplasy, the topological discordance between a gene tree and a species tree that can give the illusion of homoplasy (Dasmahapatra et al. 2012).

To further investigate this hemiplasy, we generated rooted homolog trees from the unrooted homolog trees of the carnivorous Caryophyllales dataset we already used in Fig. 2 and originally published by Walker et al. in 2017. We then identified specific conflicts with the established consensus species tree in these homologs using CAnDI (Fig. 3).

**Figure 3.**
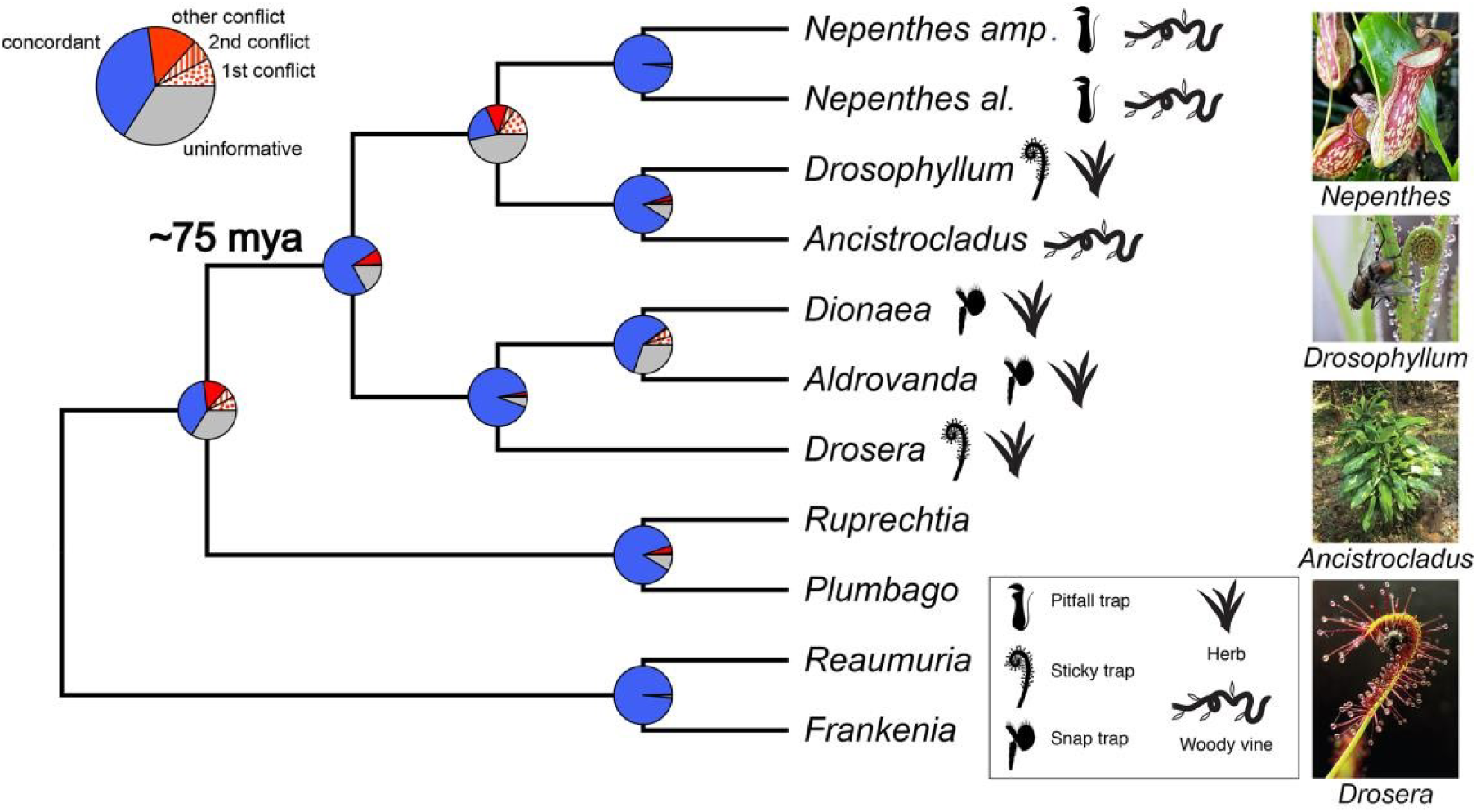
Patterns of conflict and ancestral traits are not found in all descendants (“pleisiomorphic traits”) in the carnivorous Caryophyllales. Pie charts depict the amount of gene tree conflict observed, with the blue, red and grey slices representing, respectively, the proportion of homolog gene trees concordant, conflicting and uninformative (SH-alrt ≤ 80 or missing taxon) at each node in the species tree. In case of conflict, the proportions of gene trees supporting two main alternative topologies are represented with a dotted or a striped red slice, respectively. The proportion of gene trees supporting various alternative topologies is represented with a solid red slice. The type of trap found in each carnivorous species, as well as the herbaceous or woody stem morphology, is depicted with a symbol on the right of the phylogeny. Picture credit: *Drosophyllum lusitanicum* (incidencematrix), *Ancistrocladus heyneanus* (Vinayaraj) and *Drosera capensis* (NoahElhardt) via Wikimedia Commons.

We found that an order of magnitude more conflicting gene trees support *D. lusitanicum* as sister to *Drosera binata* than the evolutionarily equivalent conflicting relationship *Ancistrocladus robertsoniorum* sister to *D. binata* (Fig. 4A). When we tested if we could recover this pattern using the orthologs inferred by the original study, we only found one ortholog tree placed *D. lusitanicum* and *D. binata* sister to one another. This pattern is only evident in homolog trees, thereby emphasising the importance of analysing as much of the phylome as possible when making biological inferences.

**Figure 4.**
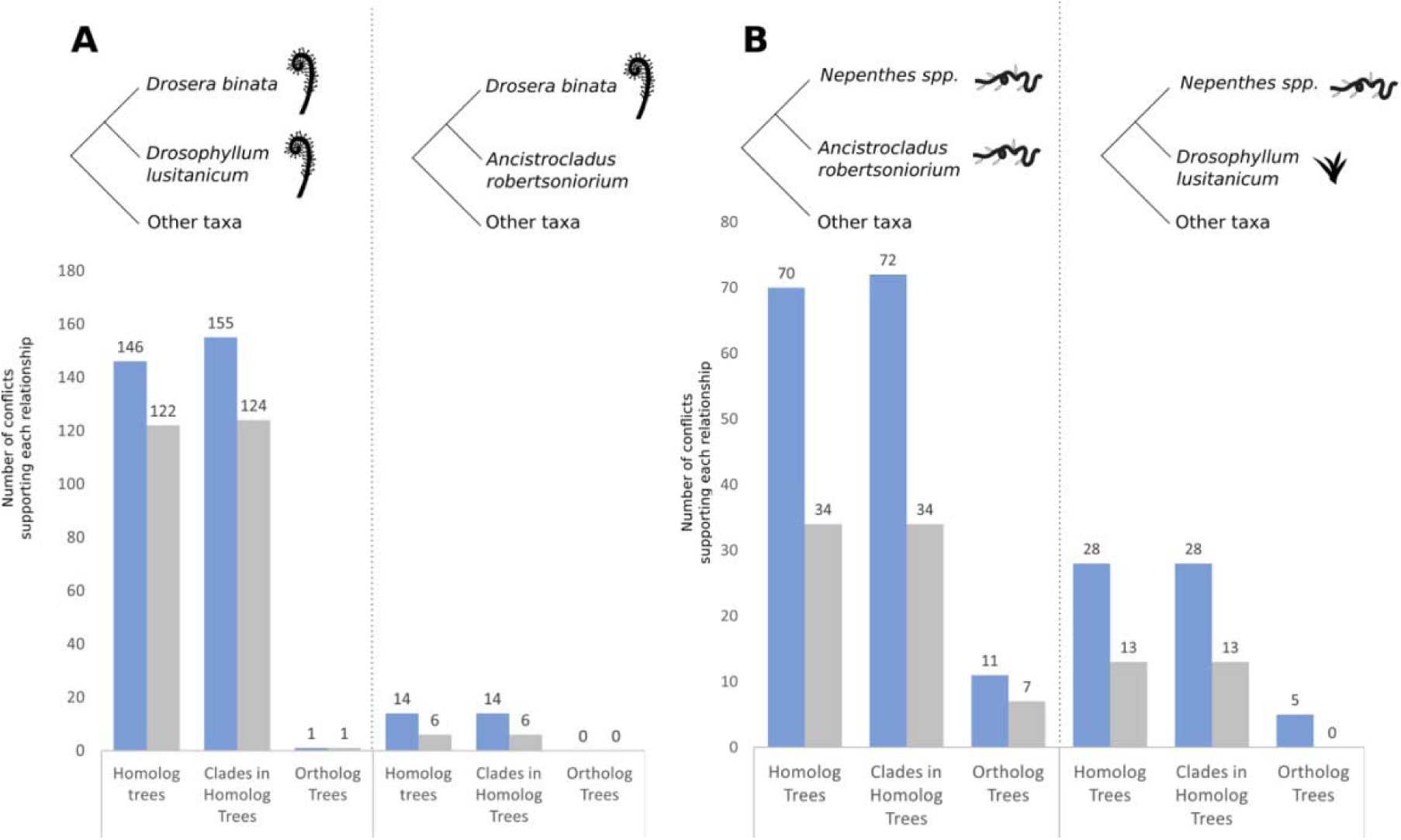
Distribution of conflicting gene trees according to the alternative species relationship they support, focusing on (A) carnivory and (B) life-history. The relationship being identified is drawn as a tree (top), while the number of conflicting gene trees supporting that relationship are shown below in homolog trees, clades within homolog trees and ortholog trees. The number of trees/subtrees for each alternative sister pair is shown in blue, followed by the number of well-supported (80 ≥ SH-alrt) trees/subtrees in grey. Homolog trees refers to the number of trees containing the relationship, while clades in homolog trees refer to every time the relationship occurred, including multiple occurrences in the same homolog tree. Conflicts that place species with shared stem and trap morphology as sister species are more common than evolutionarily equivalent conflicts that do not.

Using this dataset, we also investigated the apparent convergence of the woody vine life history phenotype in *Nepenthes* and *Ancistrocladus* (Fig. 3 and Supplementary Fig. S2). We found that, regardless of whether whole homologs or only ortholog trees were considered, more than twice as many conflicting genes placed the woody vines clades as sister to one another, compared to the less similar herbaceous Drosophyllaceae as sister to Ancistrocladaceae (Fig. 4B). Using homolog trees makes the pattern more easily detectable and robust (trend based on 98 gene trees rather than only 16), again highlighting the importance of analysing as much data as possible (Fig. 4).

It should be noted that gene trees are prone to estimation errors. Our use of homolog trees allows for the best-informed relationships, as taxon sampling is well known to increase the accuracy of phylogenetic estimation (Zwickl and Hillis 2002). Using homolog trees, which are informed by the maximum number of samples possible, we should have increased accuracy compared to extracted orthologs. To account for errors that may have arisen in the tree estimation procedure, we applied a support filter for the bipartitions, in this case requiring SH-alrt of at least 80. When the analysis was run with and without support, our results identified a biased number of genes supporting the placement of sticky traps sister to each other and woody plants sister to each other (Fig. 4). This bias, even with the filters, helps indicate this is an actual biological signal and not an artifact of a lack of information. Although some gene tree errors are inevitable, the functional association of the genes in this study provides further support for the signal and is an important consideration for any gene identification approach.

It is also worth noting that the family Dioncophyllaceae, which contains only three known species, two non-carnivorous lianas and the likely-facultatively carnivorous liana *Triphyophyllum peltatum*, is not represented in this dataset, but the literature strongly supports Dioncophyllaceae sister to Ancistrocladaceae (Baker et al. 2022). An analysis including *T. peltatum* and seeking conflicts that cluster *T. peltatum*, *Drosera* and *Drosophyllum* could provide additional information regarding life history and trap morphology evolution.

There are several reasons these homolog trees might display conflict. The probability of true molecular convergence is extremely low, given the large number of genes and sites displaying this pattern. Moreover, despite the short branch lengths separating *Drosera* and *Drosophyllum* in many gene trees (Supplementary Fig. S3), recent introgression is similarly unlikely, given the distant relationship between these species (*Drosera* and *Drosophyllum* diverged ∼75 mya). However, distinguishing ancient reticulation from ancient incomplete lineage sorting would be challenging, given the age of the relationships in question. Regardless of the mechanism of acquisition, it remains that adaptive selection on these genes is likely responsible for the retention of specific allelic variants post-acquisition.

### Genes that support Drosophyllaceae as sister to *Drosera* tend to be associated with the carnivorous syndrome

Genes whose evolutionary history conflicts with the species tree but places species with similar morphologies (i.e., trap type) sister to each other could help identify some of the molecular bases associated with the emergence of a given trait and likely to have contributed to adaptation (Lee et al. 2011). We therefore performed gene ontology (GO) term analyses on the gene trees that CAnDI identified as conflicting with the species tree, which instead placed the sundews *Drosera* and *Drosophyllum* as sister to one another. These analyses revealed that the functional categories associated with genes exhibiting this conflicting phylogenetic relationship are often related to carnivory (Tables 1 and 2). Cellular response to ozone (GO:0071457) and to reactive oxygen species (GO:0034614), and superoxide dismutase activity (GO:0004784), emerged as the most overrepresented biological process and molecular function, respectively. Carnivorous species have been shown to use free radicals during the digestion process (Chia et al. 2004), but these can also cause cellular damage. Hence, genes with enzymatic activities capable of scavenging superoxide radicals (Bowler et al. 1991) could play a key protective role in carnivorous plants. GO terms associated with nitrogen utilization (i.e., GO:0019676, GO:0019740, GO:0004356) are also among the top categories enriched in the set of genes exhibiting conflict and could relate to the assimilation of nutrients following prey digestion. Our analyses also reveal enrichment patterns for GO terms connected with photosynthesis and respiration (i.e., GO:0016168, GO:0046933, GO:0009768). Producing the morphology and molecules that allow carnivory (i.e., pigment to attract insects or traps to catch them) is energetically demanding, and carnivory impacts both respiration and photosynthesis (Adamec et al., 2021; Pavlovič, 2022). Indeed, trap development and activity are generally accompanied by spatio-temporal activation of respiration and inactivation of photosynthesis. Following this, photosynthesis can increase as a result of feeding (Pavlovič and Saganová 2015). Our analysis also points toward other interesting categories. For instance, genes connected to salicylic acid binding (GO:1901149) are overrepresented among the genes in conflict with the species tree that group the two genera of sundews as sisters to each other (Table 1). Salicylic acid (SA) and jasmonic acid (JA) are well-known to work together to regulate plant immunity (Pieterse et al., 2012). Interestingly, recent results revealed that JA signaling for plant defense has likely been co-opted by some Caryophyllales to support carnivory (Pavlovič and Saganová, 2015; Bemm et al., 2016; Pavlovič and Mithöfer, 2019; Pavlovič et al., 2024). Whether this indicates that the crosstalk between SA and JA can possibly mediate carnivory-related function remains to be established.

**Table 1.** Result of Gene Ontology analyses (Molecular Function) performed on genes in conflict with the species tree but supporting the *Drosera* + *Drosophyllum* relationship. The top ten most enriched categories from the ‘Molecular Function’ database are indicated. An extended version of this table, including subcategories, is provided in Supplementary Spreadsheet 1. FE, Fold Enrichment; FDR, False Discovery Rate.

| GO Term | Description | Expected (Observed) | FE | Raw P-value | FDR |
| --- | --- | --- | --- | --- | --- |
| GO:0004784 | Superoxide dismutase activity | 0.04(3) | 78.02 | 1.72e-05 | 1.74e-03 |
| GO:0004356 | glutamate-ammonia ligase activity | 0.03(2) | 69.35 | 6.24e-04 | 3.90e-02 |
| GO:0030060 | L-malate dehydrogenase activity | 0.05(3) | 62.41 | 2.95e-05 | 2.74e-03 |
| GO:0003746 | translation elongation factor activity | 0.08(5) | 61.19 | 5.73e-08 | 9.78e-06 |
| GO:0016168 | Chlorophyll binding | 0.13(7) | 52.01 | 2.94e-10 | 7.94e-08 |
| GO:0008266 | Poly(U) RNA binding | 0.13(6) | 46.23 | 1.06e-08 | 2.16e-06 |
| GO:1901149 | salicylic acid binding | 0.13(5) | 37.15 | 4.95e-07 | 8.03e-05 |
| GO:0046933 | proton-transporting ATP<br>synthase activity, rotational<br>mechanism | 0.12(3) | 26.01 | 2.88e-04 | 1.99e-02 |
| GO:0016762 | xyloglucan:xyloglucosyl<br>transferase activity | 0.15(3) | 19.50 | 6.26e-04 | 3.84e-02 |
| GO:0004601 | Peroxidase activity | 0.37(6) | 16.42 | 2.78e-06 | 3.61e-04 |

**Table 2.**
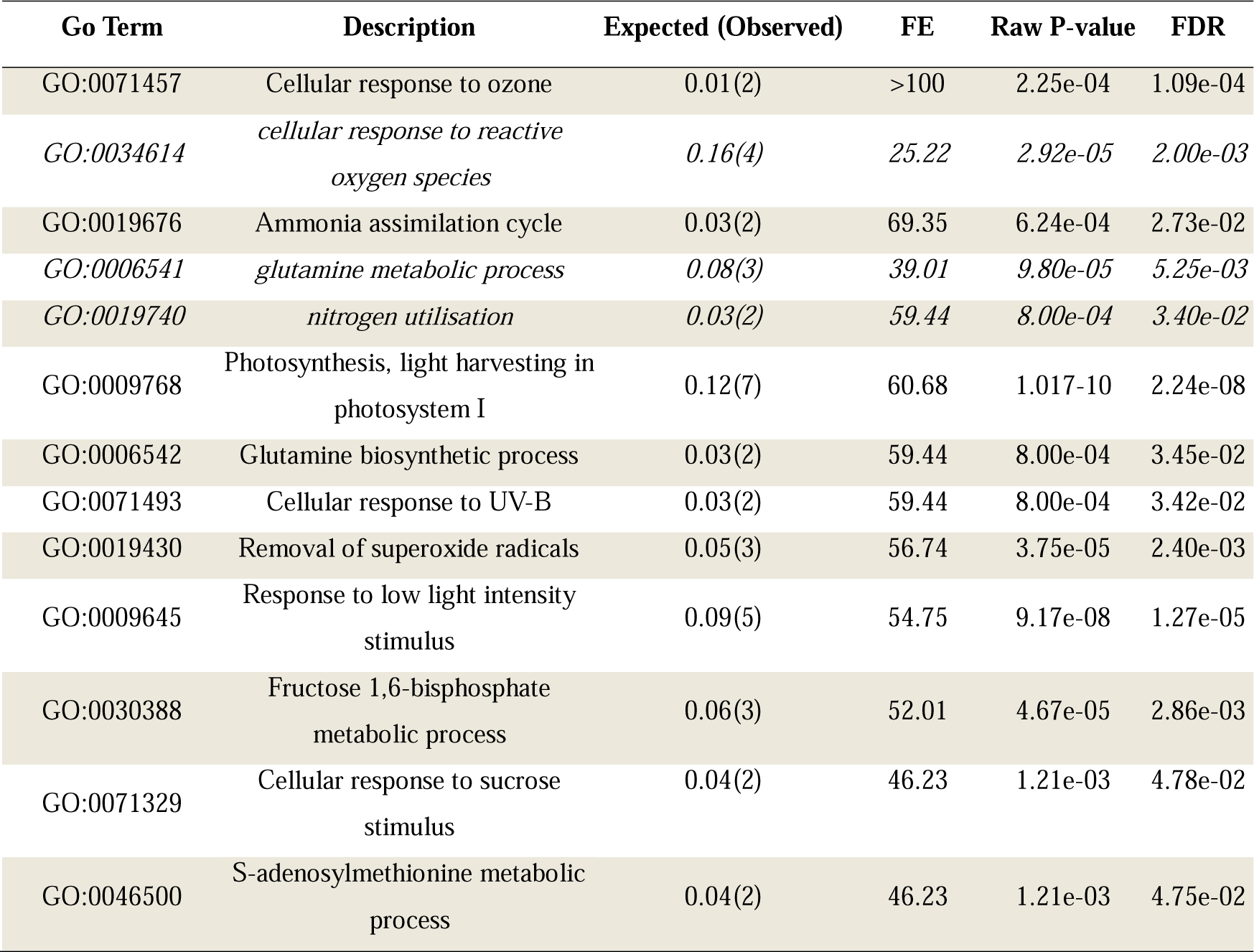
Result of Gene Ontology analyses (Biological Process) performed on genes in conflict with the species tree but supporting the *Drosera* + *Drosophyllum* sister relationship. The main categories enriched from the ‘Biological Process’ database are indicated, along with selected related categories (italicized). An extended version of this table, including all subcategories, is provided in Supplementary Spreadsheet 2. FE, Fold Enrichment; FDR, False Discovery Rate.

| Go Term | Description | Expected (Observed) | FE | Raw P-value | FDR |
| --- | --- | --- | --- | --- | --- |
| GO:0071457 | Cellular response to ozone | 0.01(2) | >100 | 2.25e-04 | 1.09e-04 |
| GO:0034614 | *cellular response to reactive oxygen species* | *0.16(4)* | *25.22* | *2.92e-05* | *2.00e-03* |
| GO:0019676 | Ammonia assimilation cycle | 0.03(2) | 69.35 | 6.24e-04 | 2.73e-02 |
| GO:0006541 | *glutamine metabolic process* | *0.08(3)* | *39.01* | *9.80e-05* | *5.25e-03* |
| GO:0019740 | *nitrogen utilisation* | *0.03(2)* | *59.44* | *8.00e-04* | *3.40e-02* |
| GO:0009768 | Photosynthesis, light harvesting in photosystem I | 0.12(7) | 60.68 | 1.017e-10 | 2.24e-08 |
| GO:0006542 | Glutamine biosynthetic process | 0.03(2) | 59.44 | 8.00e-04 | 3.45e-02 |
| GO:0071493 | Cellular response to UV-B | 0.03(2) | 59.44 | 8.00e-04 | 3.42e-02 |
| GO:0019430 | Removal of superoxide radicals | 0.05(3) | 56.74 | 3.75e-05 | 2.40e-03 |
| GO:0009645 | Response to low light intensity stimulus | 0.09(5) | 54.75 | 9.17e-08 | 1.27e-05 |
| GO:0030388 | Fructose 1,6-bisphosphate metabolic process | 0.06(3) | 52.01 | 4.67e-05 | 2.86e-03 |
| GO:0071329 | Cellular response to sucrose stimulus | 0.04(2) | 46.23 | 1.21e-03 | 4.78e-02 |
| GO:0046500 | S-adenosylmethionine metabolic process | 0.04(2) | 46.23 | 1.21e-03 | 4.75e-02 |

Next, we conducted a detailed examination of the families to which these conflicting gene trees belong. Plant carnivory is typified by a set of characters that enable them to attract, capture, digest and assimilate animal prey. This cluster of attributes is known as the carnivorous syndrome. Within the genes exhibiting conflict, we found families participating in each of the four facets of the carnivorous syndrome (Table 3 and Supplementary Fig. S3); for example, three gene clusters code for enzymes participating in anthocyanin synthesis. Red anthocyanin pigments colour the sticky trap of both *Drosera* and *Drosophyllum*. Furthermore, while the purpose of anthocyanin pigmentation in carnivorous species is debated, red-pigmented traps are widely found in nature, and pigment accumulation is unmistakably associated with carnivory (Schefer and Ruxton 2008; Jürgens et al. 2015; Gilbert et al. 2018). Sticky trap leaves and their glandular emergences (tentacles) can bend toward stuck prey to secure their capture. Such movements are generally the product of acid-growth processes (Poppinga et al. 2013). Acid-growth allows cell extension and growth following acidification of the cell wall. Three gene clusters in the set encode Xyloglucan endotransglucosylases/hydrolases, proteins capable of regulating cell wall extensibility in response to wall acidification (Majda and Robert 2018). Enzymes found in trap secretions and mediating prey digestion (i.e., chitinase) or protecting plant cells from damages caused by free radicals used during the digestion process (i.e., peroxidases) are also overrepresented among conflicting genes. Finally, genes encoding proteins contributing to the utilisation of nitrogen (for growth and development) or to the use of amino acids (alternative respiratory substrate to fuel the energy demand of carnivory) from digested prey are also present (Table 3). As *Drosera* and *Drosophyllum* resemble each other in terms of morphology and the action mechanism of their traps, these genes with histories that conflict with the species tree represent candidate genes that could have participated in the emergence of the sticky trap phenotype ∼75 Mya, or that may have mediated the parallel evolution of sticky traps in both groups of sundews.

**Table 3.**
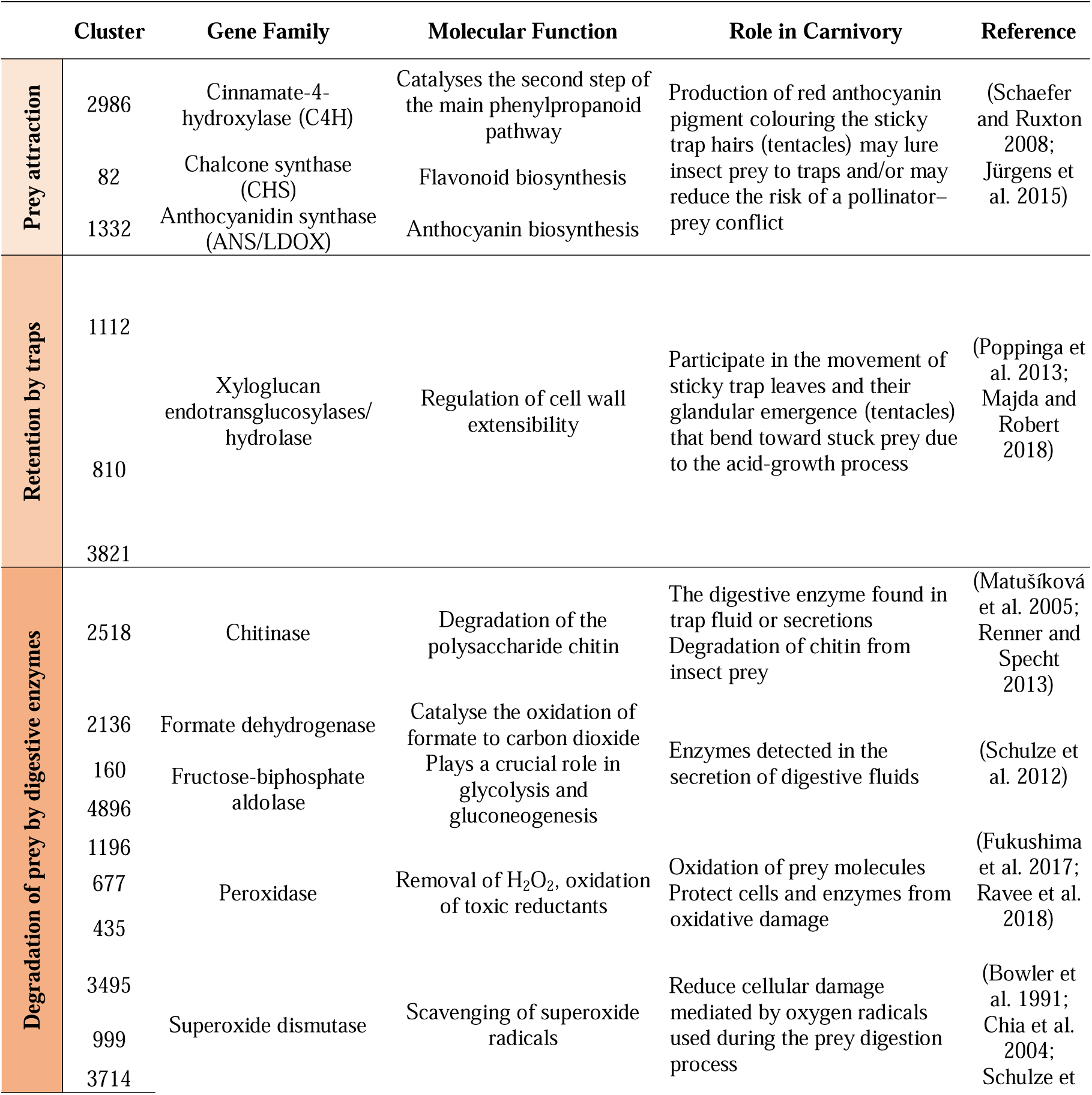
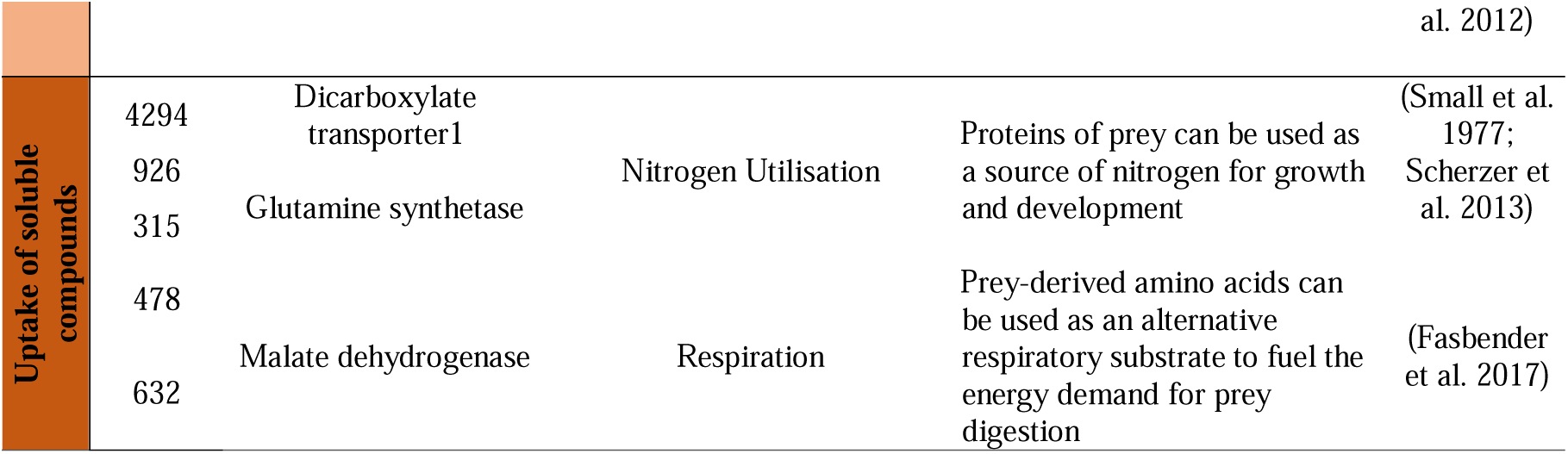
Genes in conflict with species relationships that support the relationship *Drosera* + *Drosophyllum* and encoding protein belonging to families with carnivory associated functions. The first column indicates the aspect of the carnivorous syndrome associated with selected clusters.

### An in-depth analysis of *Chitinase* (*CTL1*) reveals a history of duplication and conflict

We found a class I chitinase among the genes placing *Drosophyllum* and *Drosera* as sister to one another (Table 3). The class I chitinase gene family plays an essential role in plant evolutionary defense against fungal pathogens. Producing chitinase allows plants to break down the chitinous fungal cell wall (Samac et al. 1990; Brogue et al. 1991). Two subclasses of class I chitinases have been identified in the carnivorous Caryophyllales, with subclass Ia participating in the pathogenesis response. In contrast, members of subclass Ib help break down the chitinous exoskeleton of insects, supporting carnivory (Renner and Specht 2012).

Although a commonly performed mapping approach is gene tree to species tree, the ability to map a species tree onto a gene tree provides a powerful tool for single gene family analyses. Species tree reconciliation algorithms such as NOTUNG (Chen et al. 2000) map duplications and other events to gene trees, but CAnDI does so in homolog trees with a focus on conflict. By mapping the species tree onto the class I chitinase homolog tree (Fig. 5A), we found evidence for the previously characterised duplication of class I chitinases at the base of the carnivorous Caryophyllales (Renner and Specht 2012). Although the carnivorous Caryophyllales are noted for their high propensity of genome duplication (Walker et al. 2017; Palfalvi et al. 2020), *Drosophyllum* does not have identifiable evidence for a whole genome duplication (Walker et al. 2017). As this class I chitinase duplication is also conserved in *Drosophyllum*, it is likely not a byproduct of whole genome duplication, though it remains possible that a combination of whole genome duplication and local duplication assisted the spread of the chitinases through this clade.

**Figure 5.**
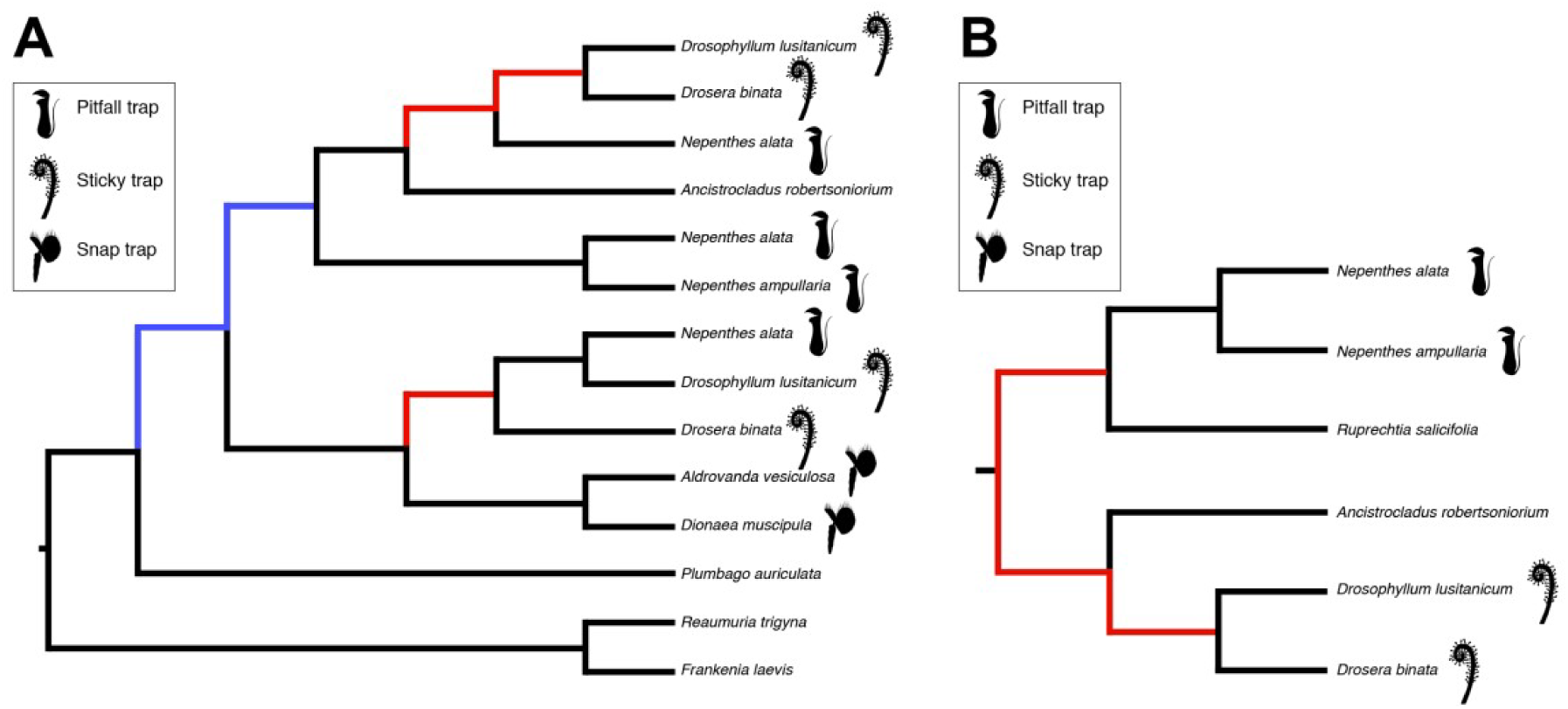
Gene duplication and conflict in the evolution of (A) the chitinase I gene (left) and (B) the *LBD4-like* gene (right). black = concordance, red = conflict, blue = duplication. The type of trap found in each carnivorous species is depicted with a symbol on the right of the homolog tree.

### An LBD transcription factor may play a key role in the development of the carnivorous trap

We first discussed the chitinase gene family due to its critical role in the evolution of plant carnivory and the focus other researchers have placed on this family in distantly related clades of carnivorous plants (Fukushima et al. 2017; Lan et al. 2017; Saul et al. 2023). However, our interest was drawn to the bias in patterns of conflict, which indicates some form of selection driving the mucilaginous trap in the carnivorous Caryophyllales. This is also suggested by the GO term analysis (Table 3). Stochastic mapping on species trees has helped researchers infer the possible ancestral states that have led to carnivory (Renner and Specht 2011). We show that complementing this work with examinations of the evolutionary history of individual gene trees can provide further insight into the evolution of plant carnivory. Such analyses can be performed on genes already known to participate in the trait of interest, such as class I chitinases, and further test previous hypotheses. Importantly, CAnDI can also single out genes with conflicting histories that have not previously been linked to carnivory, providing useful starting points to investigate their potential role.

For instance, we found one gene coding for a transcription factor among the set of 146 conflicting genes that support a sister relationship between the sundews *Drosophyllum* and *Drosera* (Fig. 5B). This transcription factor belongs to the *LATERAL ORGAN BOUNDARY DOMAIN* (*LBD*) gene family, and its closest relative in the model plant Arabidopsis is *LBD4*. It will hereafter be referred to as *LBD4-like*, since we have not determined whether this caryophyllid gene and Arabidopsis *LBD4* form a true clade. Members of the *LBD* family are regulators of plant development, controlling, for instance, leaf morphogenesis (Iwakawa et al. 2002; Shuai et al. 2002). They also fulfil important roles in regulating anthocyanin and nitrogen metabolism (Rubin et al. 2009). Interestingly, LBD homologues have also been implicated in nyctinasty (Chen et al. 2012), a behaviour characterised by pronounced leaf movements in response to the day/night cycle. In 2013, Poppinga and colleagues highlighted the need to investigate the function of *LBD* genes in other plant families also producing motor organs, including the snap trap of the carnivorous *Dionea*, in order to understand better the origins and evolution of plant lateral motor organs (Poppinga et al. 2013). Regulators of sticky trap formation should start being expressed early during leaf development in young primordia that are yet to form the morphological features characteristic of the sticky trap. To test whether the spatio-temporal expression patterns of sundews *LBD4-like* homologs were compatible with a possible role in trap elaboration, we characterised its expression in young and mature leaves of three species producing a sticky trap: *Drosera admirabilis*, *Drosera aliciae* and *Drosera coccicaulis* (Fig. 6A-C). As inflorescences were available in the *D. coccicaulis* individuals, we also measured gene expression in those tissues for comparison (Fig. 6D).

**Figure 6.**
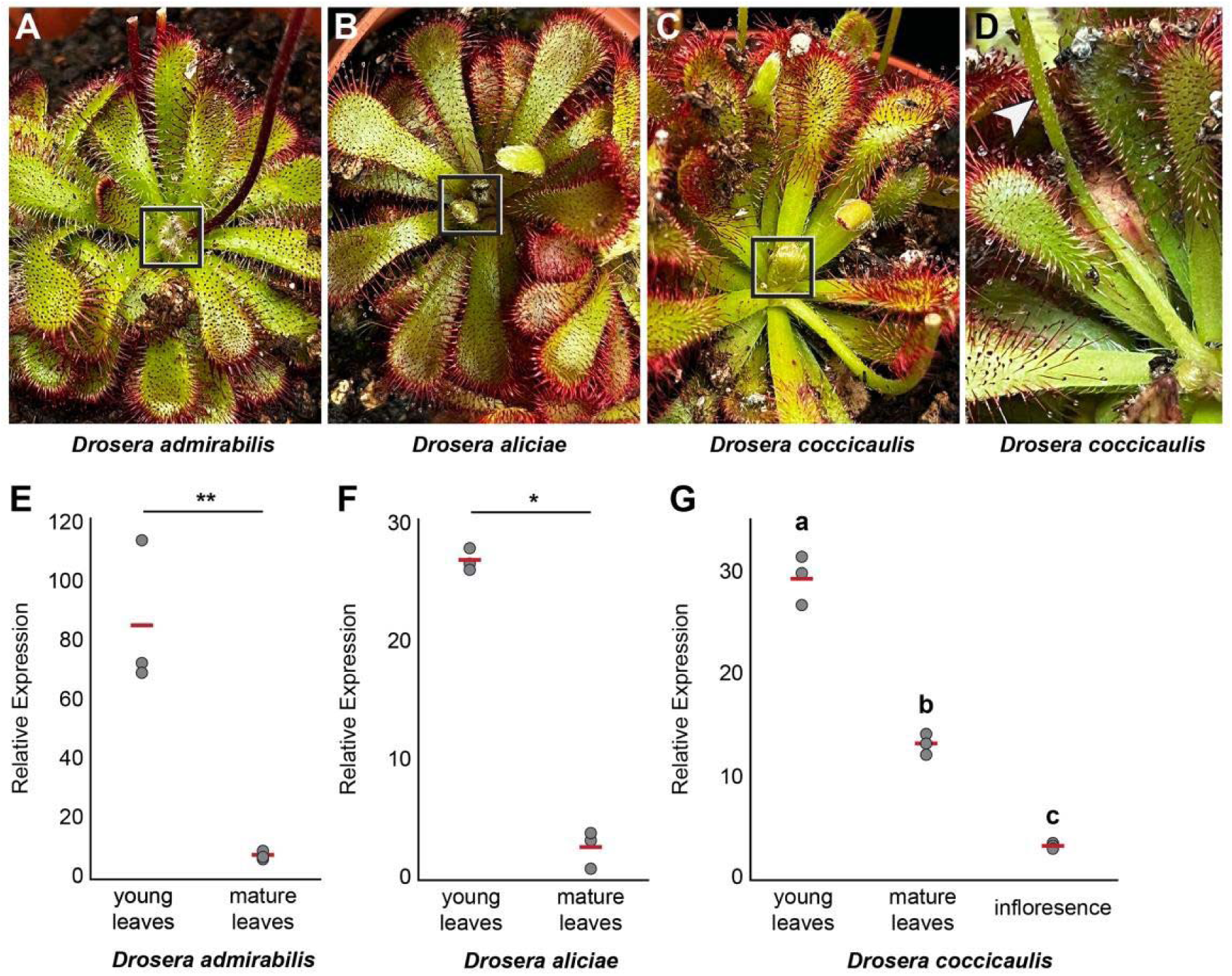
The expression dynamics of *Drosera LBD4-like* genes are compatible with a role during trap formation. **(A-D)** Young leaf primordia (boxed region) and mature traps from *Drosera admirabilis* (A), *Drosera aliciae* (B) and *Drosera coccicaulis* (C). The inflorescence of *D. coccicaulis* is partially visible in (D), indicated by a white arrowhead. **(E-F)** Relative expression levels of *LBD4-like* homologs in young leaf primordia and mature sticky traps of *D. admirabilis* (E) and *D. aliciae* (F). * p < 0.05, ** p < 0.01, Welch’s t-test. Individual dots represent distinct biological replicates (n = 3), and red lines indicate mean relative expression. (**G**) Relative expression levels of *LBD4- like* homolog in young leaf primordia, mature sticky traps and inflorescence of *D. coccicaulis*. Red lines indicate mean relative expression. Letters a, b, c, indicate the results of a post hoc Tukey’s HSD test, with group means associated with different letters being significantly different from one another (p < 0.01).

In all three species, we found that the *LBD4-like* gene was expressed in developing traps and that its expression decreases as development progresses. On average, transcript levels were two to ten times higher in young leaf primordia compared to mature ones (Fig. 6E-G). In *D. coccicaulis, LBD4-like* homolog expression was significantly weaker in reproductive tissues (inflorescence), as transcripts levels were more than eight times higher in emerging leaf primordia and almost 4 times as high in mature traps (Fig. 6G). Taken together, our results indicate that the expression dynamic of *LBD4-like* genes is compatible with a possible role in trap development in *Drosera*. Hence, *LBD4-like* is a promising candidate for functional investigation as it could be involved in shaping the unique morphology of sticky trap leaves and/or contribute to their ability to produce pigments or assimilate nitrogen following prey capture.

## Conclusion

We developed the new program CAnDI to systematically investigate patterns of gene tree conflict across the whole phylome, specifically enabling the unbiased analysis of conflict in whole homologous trees. We demonstrated that CAnDI allows for far more comparisons between gene trees and the species tree to be made using phylogenomic datasets, regardless of the identity of the organisms considered, in six previously published phylogenomic datasets. We then focused on the carnivorous Caryophyllales, showing that both carnivory and woodiness were traits associated with gene tree conflict by identifying 146 genes supporting a sister relationship between the two genera of sundews, directly conflicting with the established phylogeny of the carnivorous Caryophyllales. This supports existing evidence that phenotypic convergence and/or new adaptations frequently rely on recruiting ancestral genetic variation (Pease et al. 2016; Schweizer et al. 2019). We uncovered a clear link between those 146 genes with conflicting histories and all aspects of the carnivory syndrome. We also discovered that an *LBD4* homolog is a possible candidate regulator of sticky trap development. Hence, CAnDI enables focused examination of genes previously known to participate in a trait of interest but also provides promising means to identify novel candidates for functional investigation. This illustrates how examining patterns of gene tree conflict can further our understanding of how evolution shapes developmental processes and generates phenotypic convergences.

## Supplementary Material

The Supplementary Material file contains:

**Fig. S1** Ancestral state estimation of the trap morphology in the carnivorous Caryophyllales supports a sticky trap ancestor.

**Fig. S2** Ancestral state estimation of the growth habit in the carnivorous Caryophyllales.

**Fig. S3** Graphical representation of all the trees listed in Table 3 of the main text.

**Fig. S4** Example of examining the relationship between a homolog tree and a species tree.

**Fig. S5** Picture of young leaf and mature trap tissues used for qRT-PCR assay.

**Supplementary Method –** Ancestral state reconstruction

**Supporting References**

Other supporting materials for this manuscript include the following:

**Supplementary Spreadsheet 1.** Outcome of Gene Ontology Analysis (Molecular Function)

**Supplementary Spreadsheet 2.** Outcome of Gene Ontology Analysis (Biological Process)

**Supplementary Spreadsheet 3.** List of Primers used in this study

## Supporting information

Supplementary material_Robertson 2024

## Acknowledgments and Funding

We would like to thank Yuxi Liang for drawing the three silhouettes representing the different trap systems used in Fig. 3-5 and Fig. S1-S2. Stephen Smith for helpful discussions on the best procedure for identifying duplication events. Andrew Alverson, Erik Koenen, Diego Morales-Briones, and Drew Larson for assistance with dataset acquisition and formatting.

We are also grateful to Drew Larson, Chris Whitewoods and the members of the Moyroud lab and Walker lab for useful discussions and comments on earlier versions of this manuscript. Our research is supported by the Gatsby Charitable Foundation, the Isaac Newton Trust/Wellcome Trust ISSF, Murray Edwards College and the SBS-DTP PhD program at the University of Cambridge and start-up funding from the University of Illinois at Chicago.

## References

Adamec, L., Matušíková, I. and Pavlovič, A., 2021. Recent ecophysiological, biochemical and evolutional insights into plant carnivory. Ann. Bot., 128:241–259.

Ane C., Larget B., Baum D.A., Smith S.D., Rokas A. 2006. Bayesian Estimation of Concordance among Gene Trees. Mol. Biol. Evol. 24:412–426.

Arai N., Ohno Y., Jumyo S., Hamaji Y., Ohyama T. 2021. Organ-specific expression and epigenetic traits of genes encoding digestive enzymes in the lance-leaf sundew (Drosera adelae). J. Exp. Bot. 72:1946–1961.

Arbuckle K., Bennett C.M., Speed M.P. 2014. A simple measure of the strength of convergent evolution. Methods Ecol. Evol. 5:685–693.

Baker W.J., Bailey P., Barber V., Barker A., Bellot S., Bishop D., Botigué L.R., Brewer G., Carruthers T., Clarkson J.J., Cook J., Cowan R.S., Dodsworth S., Epitawalage N., Françoso E., Gallego B., Johnson M.G., Kim J.T., Leempoel K., Maurin O., Mcginnie C., Pokorny L., Roy S., Stone M., Toledo E., Wickett N.J., Zuntini A.R., Eiserhardt W.L., Kersey P.J., Leitch I.J., Forest F. 2022. A Comprehensive Phylogenomic Platform for Exploring the Angiosperm Tree of Life. Syst. Biol. 71:301–319.

Baum D.A. 2007. Concordance trees, concordance factors, and the exploration of reticulate genealogy. TAXON. 56:417–426.

Bemm F, Becker D, Larisch C, et al. . 2016. Venus flytrap carnivorous lifestyle builds on herbivore defence strategies. Genome Research 26, 812–825.

Bettisworth B., Stamatakis A. 2021. Root Digger: a root placement program for phylogenetic trees. BMC Bioinformatics. 22:225.

Bowler C., Slooten L., Vendenbranden S., De Rycke R., Botterman J., Sybesma C., Van Montagu marc, Inzé D. 1991. Manganese superoxide dismutase can reduce cellular damage mediated by oxygen radicals in transgenic plants. EMBO J. 10:1723–1732.

Brogue K., Chet I., Holliday M., Cressman R., Biddle P., Knowlton S., Mauvais C.J., Broglie R. 1991. Transgenic Plants with Enhanced Resistance to the Fungal Pathogen *Rhizoctonia solani*. Science. 254:1194–1197.

Brown J.M., Thomson R.C. 2016. Bayes factors unmask highly variable information content, bias, and extreme influence in phylogenomic analyses. Syst. Biol.:syw101.

Brown J.W., Walker J.F., Smith S.A. 2017. Phyx: phylogenetic tools for unix. Bioinformatics. 33:1886–1888.

Cameron K.M., Wurdack K.J., Jobson R.W. 2002. Molecular evidence for the common origin of snap-traps among carnivorous plants. Am. J. Bot. 89:1503–1509.

Chen J., Moreau C., Liu Y., Kawaguchi M., Hofer J., Ellis N., Chen R. 2012. Conserved genetic determinant of motor organ identity in Medicago truncatula and related legumes. Proc. Natl. Acad. Sci. 109:11723–11728.

Chen K., Durand D., Farach-Colton M. 2000. Notung: dating gene duplications using gene family trees. Proc. Fourth Annu. Int. Conf. Comput. Mol. Biol.:96–106.

Chia T.F., Aung H.H., Osipov A.N., Goh N.K., Chia L.S. 2004. Carnivorous pitcher plant uses free radicals in the digestion of prey. Redox Rep. 9:255–261.

Chiari Y., Cahais V., Galtier N., Delsuc F. 2012. Phylogenomic analyses support the position of turtles as the sister group of birds and crocodiles (Archosauria). BMC Biol. 10:65.

Clark J.W., Donoghue P.C.J. 2018. Whole-Genome Duplication and Plant Macroevolution. Trends Plant Sci. 23:933–945.

Copetti D., Búrquez A., Bustamante E., Charboneau J.L.M., Childs K.L., Eguiarte L.E., Lee S., Liu T.L., McMahon M.M., Whiteman N.K., Wing R.A., Wojciechowski M.F., Sanderson M.J. 2017. Extensive gene tree discordance and hemiplasy shaped the genomes of North American columnar cacti. Proc. Natl. Acad. Sci. 114:12003–12008.

Copley S.D. 2020. Evolution of new enzymes by gene duplication and divergence. FEBS J. 287:1262–1283.

Corbett-Detig R.B., Russell S.L., Nielsen R., Losos J. 2020. Phenotypic Convergence Is Not Mirrored at the Protein Level in a Lizard Adaptive Radiation. Mol. Biol. Evol. 37:1604–1614.

Dasmahapatra K.K., Walters J.R., Briscoe A.D., Davey J.W., Whibley A., Nadeau N.J., Zimin A.V., Hughes D.S.T., Ferguson L.C., Martin S.H., Salazar C., Lewis J.J., Adler S., Ahn S.-J., Baker D.A., Baxter S.W., Chamberlain N.L., Chauhan R., Counterman B.A., Dalmay T., Gilbert L.E., Gordon K., Heckel D.G., Hines H.M., Hoff K.J., Holland P.W.H., Jacquin-Joly E., Jiggins F.M., Jones R.T., Kapan D.D., Kersey P., Lamas G., Lawson D., Mapleson D., Maroja L.S., Martin A., Moxon S., Palmer W.J., Papa R., Papanicolaou A., Pauchet Y., Ray D.A., Rosser N., Salzberg S.L., Supple M.A., Surridge A., Tenger-Trolander A., Vogel H., Wilkinson P.A., Wilson D., Yorke J.A., Yuan F., Balmuth A.L., Eland C., Gharbi K., Thomson M., Gibbs R.A., Han Y., Jayaseelan J.C., Kovar C., Mathew T., Muzny D.M., Ongeri F., Pu L.-L., Qu J., Thornton R.L., Worley K.C., Wu Y.-Q., Linares M., Blaxter M.L., ffrench-Constant R.H., Joron M., Kronforst M.R., Mullen S.P., Reed R.D., Scherer S.E., Richards S., Mallet J., Owen McMillan W., Jiggins C.D., The Heliconius Genome Consortium. 2012. Butterfly genome reveals promiscuous exchange of mimicry adaptations among species. Nature. 487:94–98.

Delsuc F., Brinkmann H., Philippe H. 2005. Phylogenomics and the reconstruction of the tree of life. Nat. Rev. Genet. 6:361–375.

Doolittle R.F. 1994. Convergent evolution: the need to be explicit. Trends Biochem. Sci. 19:15– 18.

Doyle J.J. 1992. Gene Trees and Species Trees: Molecular Systematics as One-Character Taxonomy. Syst. Bot. 17:144–163.

Dunn C.W., Hejnol A., Matus D.Q., Pang K., Browne W.E., Smith S.A., Seaver E., Rouse G.W., Obst M., Edgecombe G.D., Sørensen M.V., Haddock S.H.D., Schmidt-Rhaesa A., Okusu A., Kristensen R.M., Wheeler W.C., Martindale M.Q., Giribet G. 2008. Broad phylogenomic sampling improves resolution of the animal tree of life. Nature. 452:745– 749.

Dunning L.T., Olofsson J.K., Parisod C., Choudhury R.R., Moreno-Villena J.J., Yang Y., Dionora J., Quick W.P., Park M., Bennetzen J.L., Besnard G., Nosil P., Osborne C.P., Christin P.-A. 2019. Lateral transfers of large DNA fragments spread functional genes among grasses. Proc. Natl. Acad. Sci. 116:4416–4425.

Fasbender L., Maurer D., Kreuzwieser J., Kreuzer I., Schulze W.X., Kruse J., Becker D., Alfarraj S., Hedrich R., Werner C., Rennenberg H. 2017. The carnivorous Venus flytrap uses prey-derived amino acid carbon to fuel respiration. New Phytol. 214:597–606.

Feng S., Bai M., Rivas-González I., Li C., Liu S., Tong Y., Yang H., Chen G., Xie D., Sears K.E., Franco L.M., Gaitan-Espitia J.D., Nespolo R.F., Johnson W.E., Yang H., Brandies P.A., Hogg C.J., Belov K., Renfree M.B., Helgen K.M., Boomsma J.J., Schierup M.H., Zhang G. 2022. Incomplete lineage sorting and phenotypic evolution in marsupials. Cell. 185:1646–1660.e18.

Foster C.S.P., Sauquet H., van der Merwe M., McPherson H., Rossetto M., Ho S.Y.W. 2017. Evaluating the Impact of Genomic Data and Priors on Bayesian Estimates of the Angiosperm Evolutionary Timescale. Syst. Biol. 66:338–351.

Fukushima K., Fang X., Alvarez-Ponce D., Cai H., Carretero-Paulet L., Chen C., Chang T.-H., Farr K.M., Fujita T., Hiwatashi Y., Hoshi Y., Imai T., Kasahara M., Librado P., Mao L., Mori H., Nishiyama T., Nozawa M., Pálfalvi G., Pollard S.T., Rozas J., Sánchez-Gracia A., Sankoff D., Shibata T.F., Shigenobu S., Sumikawa N., Uzawa T., Xie M., Zheng C., Pollock D.D., Albert V.A., Li S., Hasebe M. 2017. Genome of the pitcher plant Cephalotus reveals genetic changes associated with carnivory. Nat. Ecol. Evol. 1:1–9.

Ganger M.T., Dietz G.D., Ewing S.J. 2017. A common base method for analysis of qPCR data and the application of simple blocking in qPCR experiments. BMC Bioinformatics. 18:534.

Gatesy J., Baker R.H. 2005. Hidden Likelihood Support in Genomic Data: Can Forty-Five Wrongs Make a Right? Syst. Biol. 54:483–492.

Gilbert K.J., Nitta J.H., Talavera G., Pierce N.E., 2018. Keeping an eye on coloration: ecological correlates of the evolution of pitcher traits in the genus *Nepenthes* (Caryophyllales), Biol. J. Linn. Soc. 123:321–337

Hejnol A., Obst M., Stamatakis A., Ott M., Rouse G.W., Edgecombe G.D., Martinez P., Baguñà J., Bailly X., Jondelius U., Wiens M., Müller W.E.G., Seaver E., Wheeler W.C., Martindale M.Q., Giribet G., Dunn C.W. 2009. Assessing the root of bilaterian animals with scalable phylogenomic methods. Proc. R. Soc. B Biol. Sci. 276:4261–4270.

Heubl G., Bringmann G., Meimberg H. 2006. Molecular Phylogeny and Character Evolution of Carnivorous Plant Families in Caryophyllales - Revisited. Plant Biol.:821–830.

Huerta-Cepas J., Dopazo H., Dopazo J., Gabaldón T. 2007. The human phylome. Genome Biol. 8:R109.

Huerta-Cepas J., Serra F., Bork P. 2016. ETE 3: Reconstruction, Analysis, and Visualization of Phylogenomic Data. Mol. Biol. Evol. 33:1635–1638.

Iñiguez L.P., Hernández G. 2017. The Evolutionary Relationship between Alternative Splicing and Gene Duplication. Front. Genet. 8.

Iwakawa H., Ueno Y., Semiarti E., Onouchi H., Kojima S., Tsukaya H., Hasebe M., Soma T., Ikezaki M., Machida C., Machida Y. 2002. The ASYMMETRIC LEAVES2 Gene of Arabidopsis thaliana, Required for Formation of a Symmetric Flat Leaf Lamina, Encodes a Member of a Novel Family of Proteins Characterized by Cysteine Repeats and a Leucine Zipper. Plant Cell Physiol. 43:467–478.

Jeffroy O., Brinkmann H., Delsuc F., Philippe H. 2006. Phylogenomics: the beginning of incongruence? Trends Genet. 22:225–231.

Johnson B.R., Borowiec M.L., Chiu J.C., Lee E.K., Atallah J., Ward P.S. 2013. Phylogenomics Resolves Evolutionary Relationships among Ants, Bees, and Wasps. Curr. Biol. 23:2058– 2062.

Jürgens A., Witt T., Sciligo A., El-Sayed A.M. 2015. The effect of trap colour and trap-flower distance on prey and pollinator capture in carnivorous Drosera species. Funct. Ecol. 29:1026– 1037.

Kalyaanamoorthy S., Minh B.Q., Wong T.K.F., von Haeseler A., Jermiin L.S. 2017. ModelFinder: fast model selection for accurate phylogenetic estimates. Nat. Methods. 14:587– 589.

Katoh K., Standley D.M. 2013. MAFFT Multiple Sequence Alignment Software Version 7: Improvements in Performance and Usability. Mol. Biol. Evol. 30:772–780.

Koenen E.J.M., Ojeda D.I., Steeves R., Migliore J., Bakker F.T., Wieringa J.J., Kidner C., Hardy O.J., Pennington R.T., Bruneau A., Hughes C.E. 2020. Large-scale genomic sequence data resolve the deepest divergences in the legume phylogeny and support a nearsimultaneous evolutionary origin of all six subfamilies. New Phytol. 225:1355–1369.

Kozlov A.M., Darriba D., Flouri T., Morel B., Stamatakis A. 2019. RAxML-NG: a fast, scalable and user-friendly tool for maximum likelihood phylogenetic inference. Bioinformatics. 35:4453– 4455.

Lan T., Renner T., Ibarra-Laclette E., Farr K.M., Chang T.-H., Cervantes-Pérez S.A., Zheng C., Sankoff D., Tang H., Purbojati R.W., Putra A., Drautz-Moses D.I., Schuster S.C., Herrera-Estrella L., Albert V.A. 2017. Long-read sequencing uncovers the adaptive topography of a carnivorous plant genome. Proc. Natl. Acad. Sci. 114:E4435–E4441.

Larson D.A., Walker J.F., Vargas O.M., Smith S.A. 2020. A consensus phylogenomic approach highlights paleopolyploid and rapid radiation in the history of Ericales. Am. J. Bot. 107:773–789.

Lee E.K., Cibrian-Jaramillo A., Kolokotronis S.-O., Katari M.S., Stamatakis A., Ott M., Chiu J.C., Little D.P., Stevenson D.W., McCombie W.R., Martienssen R.A., Coruzzi G., DeSalle R. 2011. A Functional Phylogenomic View of the Seed Plants. PLOS Genet. 7:e1002411.

Lynch M., Force A. 2000. The Probability of Duplicate Gene Preservation by Subfunctionalization. Genetics. 154:459–473.

Maddison W.P. 1997. Gene Trees in Species Trees. Syst. Biol. 46:523–536.

Maddison W.P., Maddison D.R. 2021. Mesquite: a modular system for evolutionary analysis.

Majda M., Robert S. 2018. The Role of Auxin in Cell Wall Expansion. Int. J. Mol. Sci. 19:951.

Marcet-Houben M., Gabaldón T. 2011. TreeKO: a duplication-aware algorithm for the comparison of phylogenetic trees. Nucleic Acids Res. 39:e66.

Matušíková I., Salaj J., Moravčíková J., Mlynárová L., Nap J.-P., Libantová J. 2005. Tentacles of in vitro-grown round-leaf sundew (Drosera rotundifoliaL.) show induction of chitinase activity upon mimicking the presence of prey. Planta. 222:1020–1027.

Minh B.Q., Hahn M.W., Lanfear R. 2020. New Methods to Calculate Concordance Factors for Phylogenomic Datasets. Mol. Biol. Evol. 37:2727–2733.

Morales-Briones D.F., Kadereit G., Tefarikis D.T., Moore M.J., Smith S.A., Brockington S.F., Timoneda A., Yim W.C., Cushman J.C., Yang Y. 2021. Disentangling Sources of Gene Tree Discordance in Phylogenomic Data Sets: Testing Ancient Hybridizations in Amaranthaceae s.l. Syst. Biol. 70:219–235.

Naser-Khdour S., Minh B.Q., Lanfear R. 2021. Assessing Confidence in Root Placement on Phylogenies: An Empirical Study Using Non-Reversible Models for Mammals. 2020.07.31.230144.

Nguyen L.-T., Schmidt H.A., von Haeseler A., Minh B.Q. 2015. IQ-TREE: A Fast and Effective Stochastic Algorithm for Estimating Maximum-Likelihood Phylogenies. Mol. Biol. Evol. 32:268–274.

Palfalvi G., Hackl T., Terhoeven N., Shibata T.F., Nishiyama T., Ankenbrand M., Becker D., Förster F., Freund M., Iosip A., Kreuzer I., Saul F., Kamida C., Fukushima K., Shigenobu S., Tamada Y., Adamec L., Hoshi Y., Ueda K., Winkelmann T., Fuchs J., Schubert I., Schwacke R., Al-Rasheid K., Schultz J., Hasebe M., Hedrich R. 2020. Genomes of the Venus Flytrap and Close Relatives Unveil the Roots of Plant Carnivory. Curr. Biol. 30:2312–2320.e5.

Parins-Fukuchi C., Stull G.W., Smith S.A. 2021. Phylogenomic conflict coincides with rapid morphological innovation. Proc. Natl. Acad. Sci. 118:e2023058118.

Parks M.B., Nakov T., Ruck E.C., Wickett N.J., Alverson A.J. 2018. Phylogenomics reveals an extensive history of genome duplication in diatoms (Bacillariophyta). Am. J. Bot. 105:330–347.

Pavlovič, A., 2022. Photosynthesis in Carnivorous Plants: From Genes to Gas Exchange of Green Hunters. Critical Reviews in Plant Sciences, 41:305–320.

Pavlovič, A., Koller, J., Vrobel, O., Chamrád, I., Lenobel, R. and Tarkowski, P., 2024. Is the co-option of jasmonate signalling for botanical carnivory a universal trait for all carnivorous plants? Journal of Experimental Botany, 75:334–349.

Pavlovič A, Mithöfer A. 2019. Jasmonate signalling in carnivorous plants: copycat of plant defence mechanisms. Journal of Experimental Botany 70:3379–3389.

Pavlovič A., Saganová M. 2015. A novel insight into the cost–benefit model for the evolution of botanical carnivory. Ann. Bot. 115:1075–1092.

Pease J.B., Brown J.W., Walker J.F., Hinchliff C.E., Smith S.A. 2018. Quartet Sampling distinguishes lack of support from conflicting support in the green plant tree of life. Am. J. Bot. 105:385–403.

Pease J.B., Haak D.C., Hahn M.W., Moyle L.C. 2016. Phylogenomics Reveals Three Sources of Adaptive Variation during a Rapid Radiation. PLOS Biol. 14:e1002379.

Pieterse, C. M. J., Van der Does, D., Zamioudis, C., Leon-Reyes, A., and Van Wees, S. C. M. (2012). Hormonal modulation of plant immunity. Annu. Rev. Cell Dev. Biol. 28, 489–521.

Poppinga S., Masselter T., Speck T. 2013. Faster than their prey: New insights into the rapid movements of active carnivorous plants traps. BioEssays. 35:649–657.

R Core Team. 2017. R: A language and environment for statistical computing. R Found. Stat. Comput. Vienna Austria.

Ravee R., Salleh F. ‘Imadi M., Goh H.-H. 2018. Discovery of digestive enzymes in carnivorous plants with focus on proteases. PeerJ. 6:e4914.

Renner T., Specht C.D. 2011. A Sticky Situation: Assessing Adaptations for Plant Carnivory in the Caryophyllales by Means of Stochastic Character Mapping. Int. J. Plant Sci. 172:889–901.

Renner T., Specht C.D. 2012. Molecular and Functional Evolution of Class I Chitinases for Plant Carnivory in the Caryophyllales. Mol. Biol. Evol. 29:2971–2985.

Renner T., Specht C.D. 2013. Inside the trap: gland morphologies, digestive enzymes, and the evolution of plant carnivory in the Caryophyllales. Curr. Opin. Plant Biol. 16:436–442.

Roberts W.R., Ruck E.C., Downey K.M., Pinseel E., Alverson A.J. 2023. Resolving marine– freshwater transitions by diatoms through a fog of discordant gene trees. :2022.08.12.503770.

Robinson T.J., Ruiz-Herrera A., Avise J.C. 2008. Hemiplasy and homoplasy in the karyotypic phylogenies of mammals. Proc. Natl. Acad. Sci. 105:14477–14481.

Rokas A., Williams B.L., King N., Carroll S.B. 2003. Genome-scale approaches to resolving incongruence in molecular phylogenies. Nature. 425:798–804.

Rubin G., Tohge T., Matsuda F., Saito K., Scheible W.-R. 2009. Members of the LBD Family of Transcription Factors Repress Anthocyanin Synthesis and Affect Additional Nitrogen Responses in Arabidopsis. Plant Cell. 21:3567–3584.

Samac D.A., Hironaka C.M., Yallaly P.E., Shah D.M. 1990. Isolation and Characterization of themGenes Encoding Basic and Acidic Chitinase in Arabidopsis thaliana. Plant Physiol. 93:907– 914.

Saul, F., Scharmann, M., Wakatake, T., Rajaraman S., Marques A., Freund M., Bringmann G., Channon L., Becker D., Carroll E., Low Y.W., Lindqvist C., Gilbert K.J., Renner T., Masuda S., Richter M., Vogg G., Shirasu K., Michael T.P., Hedrich R., Albert V.A., Fukushima K., 2023. Subgenome dominance shapes novel gene evolution in the decaploid pitcher plant Nepenthes gracilis. Nat. Plants 9:2000–2015.

Schaefer H.M., Ruxton G.D. 2008. Fatal attraction: carnivorous plants roll out the red carpet to lure insects. Biol. Lett. 4:153–155.

Scherzer S., Krol E., Kreuzer I., Kruse J., Karl F., von Rüden M., Escalante-Perez M., Müller T., Rennenberg H., Al-Rasheid K.A.S., Neher E., Hedrich R. 2013. The Dionaea muscipula Ammonium Channel DmAMT1 Provides NH4+ Uptake Associated with Venus Flytrap’s Prey Digestion. Curr. Biol. 23:1649–1657.

Schulze W.X., Sanggaard K.W., Kreuzer I., Knudsen A.D., Bemm F., Thøgersen I.B., Bräutigam A., Thomsen L.R., Schliesky S., Dyrlund T.F., Escalante-Perez M., Becker D., Schultz J., Karring H., Weber A., Højrup P., Hedrich R., Enghild J.J. 2012. The Protein Composition of the Digestive Fluid from the Venus Flytrap Sheds Light on Prey Digestion Mechanisms. Mol. Cell. Proteomics. 11:1306–1319.

Schweizer M., Warmuth V., Alaei Kakhki N., Aliabadian M., Förschler M., Shirihai H., Suh A., Burri R. 2019. Parallel plumage colour evolution and introgressive hybridization in wheatears. J. Evol. Biol. 32:100–110.

Shen X.-X., Hittinger C.T., Rokas A. 2017. Contentious relationships in phylogenomic studies can be driven by a handful of genes. Nat. Ecol. Evol. 1:1–10.

Shuai B., Reynaga-Peña C.G., Springer P.S. 2002. The Lateral Organ Boundaries Gene Defines a Novel, Plant-Specific Gene Family. Plant Physiol. 129:747–761.

Sicheritz-Pontén T., Andersson S.G.E. 2001. A phylogenomic approach to microbial evolution. Nucleic Acids Res. 29:545–552.

Small G.C., Onraët A., Grierson D.S., Reynolds G. 1977. Studies on Insect-Free Growth, Development and Nitrate-Assimilating Enzymes of Drosera Aliciae Hamet. New Phytol. 79:127– 133.

Smith M.L., Hahn M.W. 2021. New Approaches for Inferring Phylogenies in the Presence of Paralogs. Trends Genet. 37:174–187.

Smith S.A., Moore M.J., Brown J.W., Yang Y. 2015. Analysis of phylogenomic datasets reveals conflict, concordance, and gene duplications with examples from animals and plants. BMC Evol. Biol. 15:150.

Smith S.A., Walker J.F. 2019. PyPHLAWD: A python tool for phylogenetic dataset construction. Methods Ecol. Evol. 10:104–108.

Stern D.L. 2013. The genetic causes of convergent evolution. Nat. Rev. Genet. 14:751–764.

Stull G.W., Qu X.-J., Parins-Fukuchi C., Yang Y.-Y., Yang J.-B., Yang Z.-Y., Hu Y., Ma H., Soltis P.S., Soltis D.E., Li D.-Z., Smith S.A., Yi T.-S. 2021. Gene duplications and phylogenomic conflict underlie major pulses of phenotypic evolution in gymnosperms. Nat. Plants. 7:1015– 1025.

Walker J.F., Yang Y., Feng T., Timoneda A., Mikenas J., Hutchison V., Edwards C., Wang N., Ahluwalia S., Olivieri J., Walker-Hale N., Majure L.C., Puente R., Kadereit G., Lauterbach M., Eggli U., Flores-Olvera H., Ochoterena H., Brockington S.F., Moore M.J., Smith S.A. 2018. From cacti to carnivores: Improved phylotranscriptomic sampling and hierarchical homology inference provide further insight into the evolution of Caryophyllales. Am. J. Bot. 105:446–462.

Walker J.F., Yang Y., Moore M.J., Mikenas J., Timoneda A., Brockington S.F., Smith S.A. 2017. Widespread paleopolyploidy, gene tree conflict, and recalcitrant relationships among the carnivorous Caryophyllales. Am. J. Bot. 104:858–867.

Wickett N.J., Mirarab S., Nguyen N., Warnow T., Carpenter E., Matasci N., Ayyampalayam S., Barker M.S., Burleigh J.G., Gitzendanner M.A., Ruhfel B.R., Wafula E., Der J.P., Graham S.W., Mathews S., Melkonian M., Soltis D.E., Soltis P.S., Miles N.W., Rothfels C.J., Pokorny L., Shaw A.J., DeGironimo L., Stevenson D.W., Surek B., Villarreal J.C., Roure B., Philippe H., dePamphilis C.W., Chen T., Deyholos M.K., Baucom R.S., Kutchan T.M., Augustin M.M., Wang J., Zhang Y., Tian Z., Yan Z., Wu X., Sun X., Wong G.K.-S., Leebens-Mack J. 2014. Phylotranscriptomic analysis of the origin and early diversification of land plants. Proc. Natl. Acad. Sci. 111:E4859–E4868.

Winkelmann T., Bringmann G., Herwig A., Hedrich R. 2023. Carnivory on demand: phosphorus deficiency induces glandular leaves in the African liana Triphyophyllum peltatum. New Phytol. 239:1140–1152.

Yang Y., Moore M.J., Brockington S.F., Soltis D.E., Wong G.K.-S., Carpenter E.J., Zhang Y., Chen L., Yan Z., Xie Y., Sage R.F., Covshoff S., Hibberd J.M., Nelson M.N., Smith S.A. 2015. Dissecting Molecular Evolution in the Highly Diverse Plant Clade Caryophyllales Using Transcriptome Sequencing. Mol. Biol. Evol. 32:2001–2014.

Yang Y., Smith S.A. 2014. Orthology Inference in Nonmodel Organisms Using Transcriptomes and Low-Coverage Genomes: Improving Accuracy and Matrix Occupancy for Phylogenomics. Mol. Biol. Evol. 31:3081–3092.

Yao G., Jin J.-J., Li H.-T., Yang J.-B., Mandala V.S., Croley M., Mostow R., Douglas N.A., Chase M.W., Christenhusz M.J.M., Soltis D.E., Soltis P.S., Smith S.A., Brockington S.F., Moore M.J., Yi T.-S., Li D.-Z. 2019. Plastid phylogenomic insights into the evolution of Caryophyllales. Mol. Phylogenet. Evol. 134:74–86.

Yap V.B., Speed T. 2005. Rooting a phylogenetic tree with nonreversible substitution models. BMC Evol. Biol. 5:2.

Zhang R., Fu X., Zhao C., Cheng J., Liao H., Wang P., Yao X., Duan X., Yuan Y., Xu G., Kramer E.M., Shan H., Kong H. 2020. Identification of the Key Regulatory Genes Involved in Elaborate Petal Development and Specialized Character Formation in Nigella damascena (Ranunculaceae). Plant Cell. 32:3095–3112.

Zhou X., Lutteropp S., Czech L., Stamatakis A., Looz M.V., Rokas A. 2020. Quartet-Based Computations of Internode Certainty Provide Robust Measures of Phylogenetic Incongruence. Syst. Biol. 69:308–324.

Zmasek C.M., Eddy S.R. 2002. RIO: Analyzing proteomes by automated phylogenomics using resampled inference of orthologs. BMC Bioinformatics. 3:14.

Zwickl D.J., Hillis D.M. 2002. Increased Taxon Sampling Greatly Reduces Phylogenetic Error. Syst. Biol. 51:588–598.

