## Supplementary material_Robertson 2024 for "CAnDI: a new tool to investigate conflict in homologous gene trees and explain convergent trait evolution"

**This PDF file includes:**

**Fig. S1** Ancestral state estimation of the trap morphology in the carnivorous Caryophyllales supports a sticky trap ancestor.

**Fig. S2** Ancestral state estimation of the growth habit in the carnivorous Caryophyllales.

**Fig. S3** Graphical representation of all the trees listed in Table 3 of the main text.

**Fig. S4** Example of examining the relationship between a homolog tree and a species tree.

**Fig. S5** Picture of young leaf and mature trap tissues used for qRT-PCR assay.

**Supplementary Method –** Ancestral state reconstruction

**Supporting References**

**Other supporting materials for this manuscript include the following:**

**Supplementary Spreadsheet 1.** Outcome of Gene Ontology Analysis (Molecular

Function)

**Supplementary Spreadsheet 2.** Outcome of Gene Ontology Analysis (Biological

Process)

**Supplementary Spreadsheet 3.** List of Primers used in this study

**Figure S1 Ancestral state estimation of the trap morphology in the carnivorous Caryophyllales supports a sticky trap ancestor.** Maximum parsimony ancestral state reconstruction was carried out on the carnivorous Caryophyllales. The placement of the carnivorous Caryophyllales within the order Caryophyllales as a whole (Heubl et al. 2006) is shown on the lower left. The state of the most recent common ancestor of all carnivorous Caryophyllales was inferred to be a sticky trap. Families Drosophyllaceae

(containing only one species, the Portuguese Sundew *D. lusitanicum*), Dioncophyllaceae

[containing *Triphyophyllum peltatum* (green) and two non-carnivorous species],

Ancistrocladaceae, Nepenthaceae (the pitcher plants) and Droseraceae are labelled, with

Droseraceae divided into the sundews (genus *Drosera*) and the snap traps (genera

*Dionaea* and *Aldrovanda*).

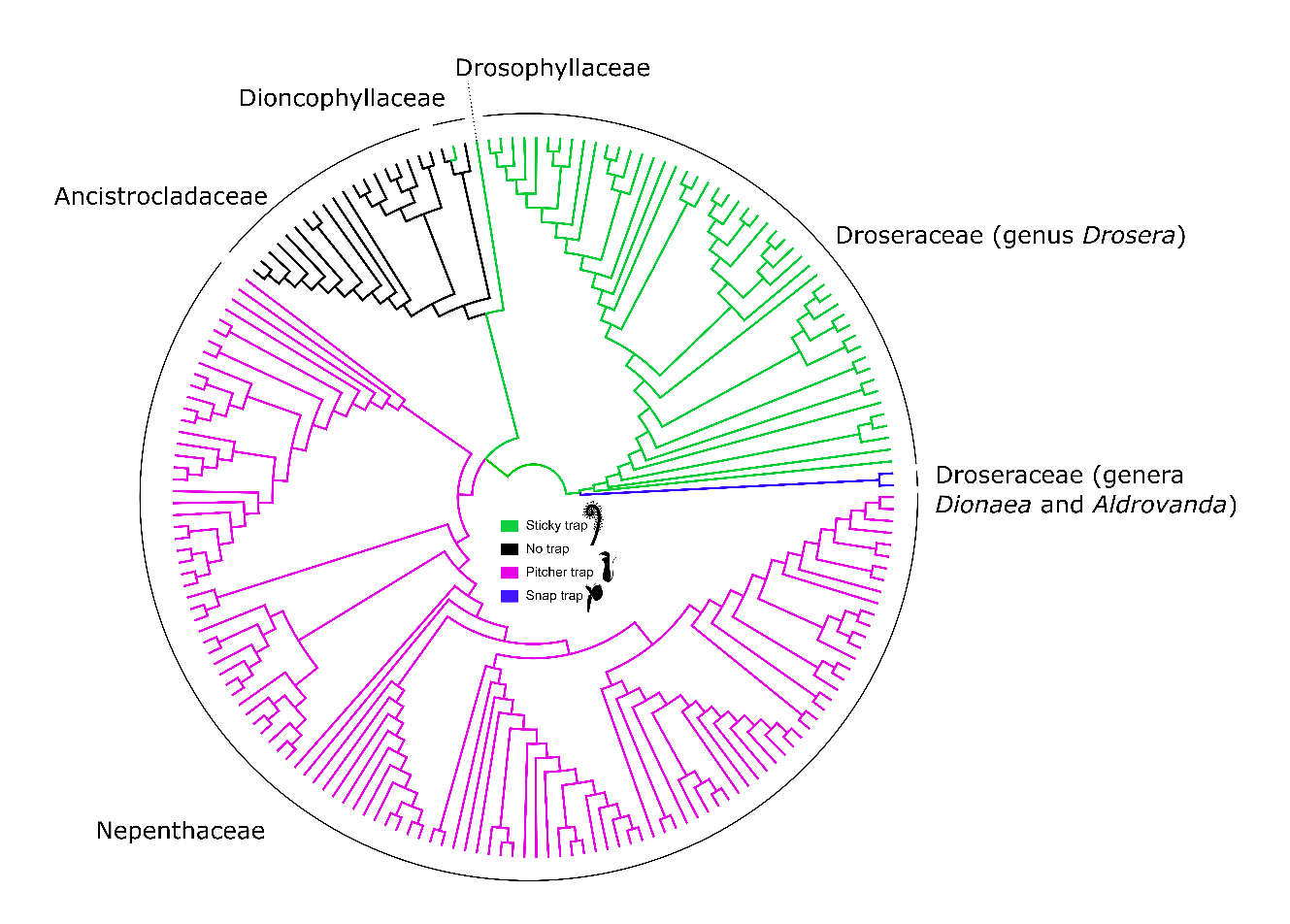

**Figure S2 Ancestral state estimation of the growth habit in the carnivorous Caryophyllales**.

Maximum parsimony ancestral state reconstruction was carried out on the carnivorous

Caryophyllales comparing woody (liana) vs herbaceous growth habits. Families Drosophyllaceae

(containing only one species, the Portuguese Sundew *D. lusitanicum*), Dioncophyllaceae

(containing *Triphyophyllum peltatum* (green) and two non-carnivorous species),

Ancistrocladaceae, Nepenthaceae (the pitcher plants) and Droseraceae are labelled, with

Droseraceae divided into the sundews (genus *Drosera*) and the snap traps (genera *Dionaea* and *Aldrovanda*). Tree was drawn using FigTree [(http://tree.bio.ed.ac.uk/software/figtree/)](http://tree.bio.ed.ac.uk/software/figtree/).

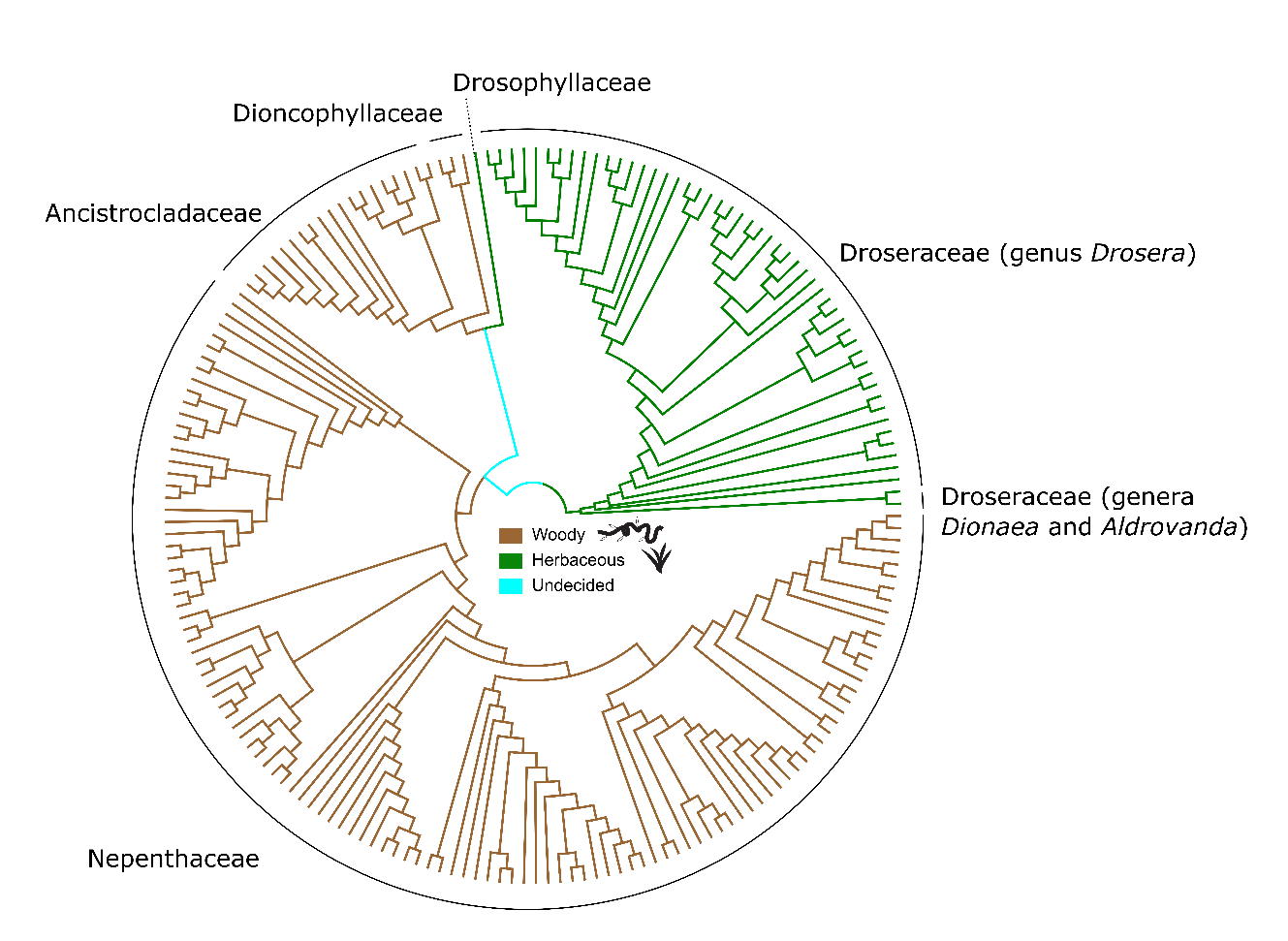

**Figure S3** **Graphical representation of all the trees listed in Table 3 of the main text.** Support (SH-aLRT) is shown at each node, and each figure contains a scale bar for branch length in substitutions per base pair. The trees are from the CARN dataset discussed in the main text. In each tree, the key *Drosera* (DrobinSFB) *x Drosophyllum* (DrolusSFB) node(s) is highlighted in blue. All trees were visualised for this figure in FigTree [(http://tree.bio.ed.ac.uk/software/figtree/)](http://tree.bio.ed.ac.uk/software/figtree/)

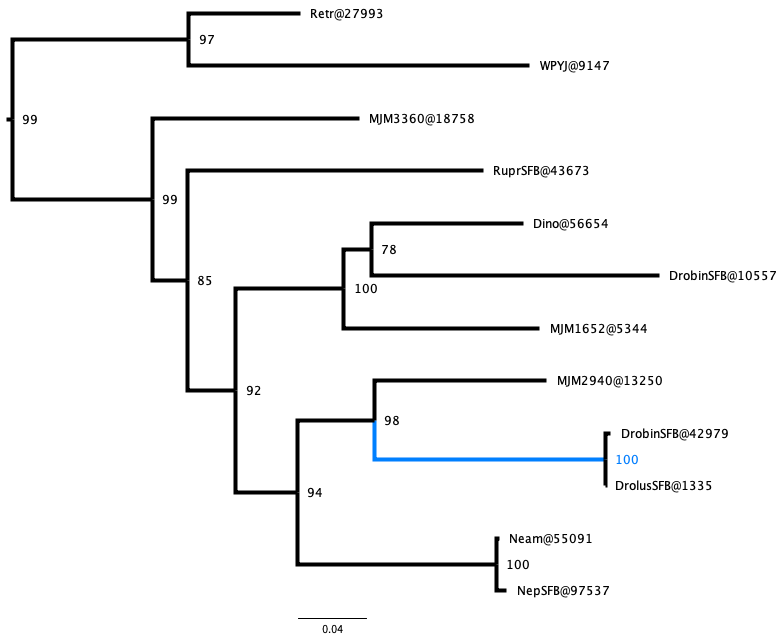

### Cinnamate-4-hydroxylase (C4H) (cluster 2986)

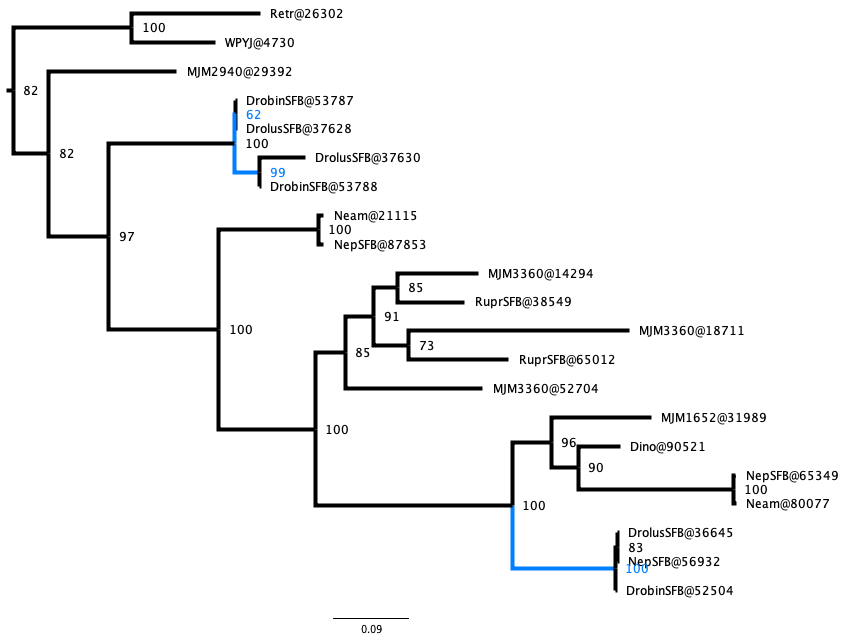

### Chalcone synthase (CHS) (cluster 82)

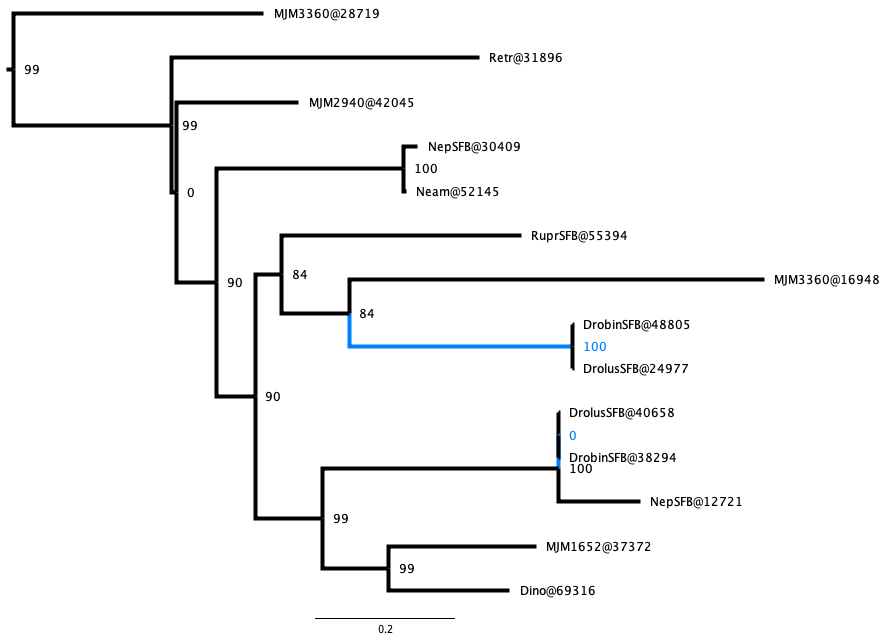

### Anthocyanidin synthase (ANS/LDOX) (cluster 1332)

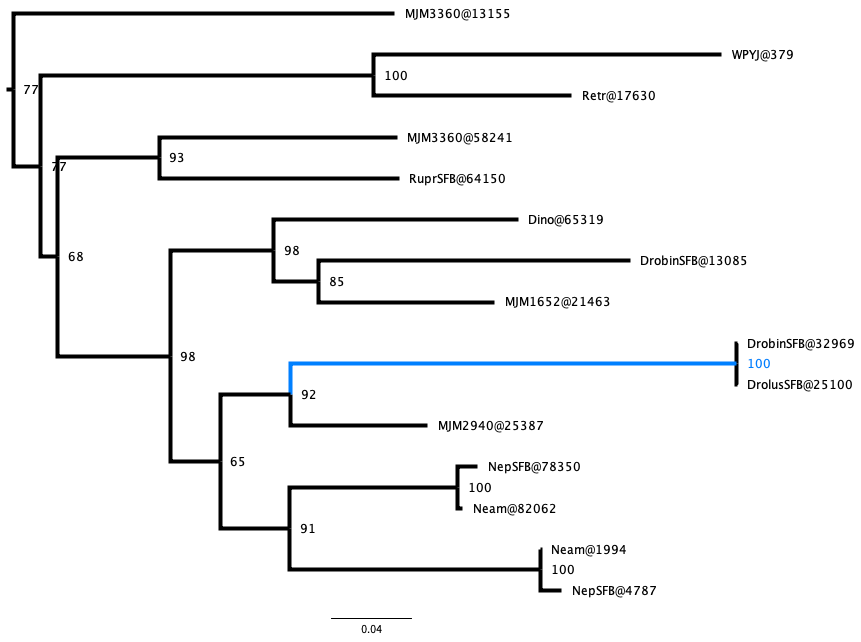

### Xyloglucan endotransglucosylases/hydrolase (cluster 810)

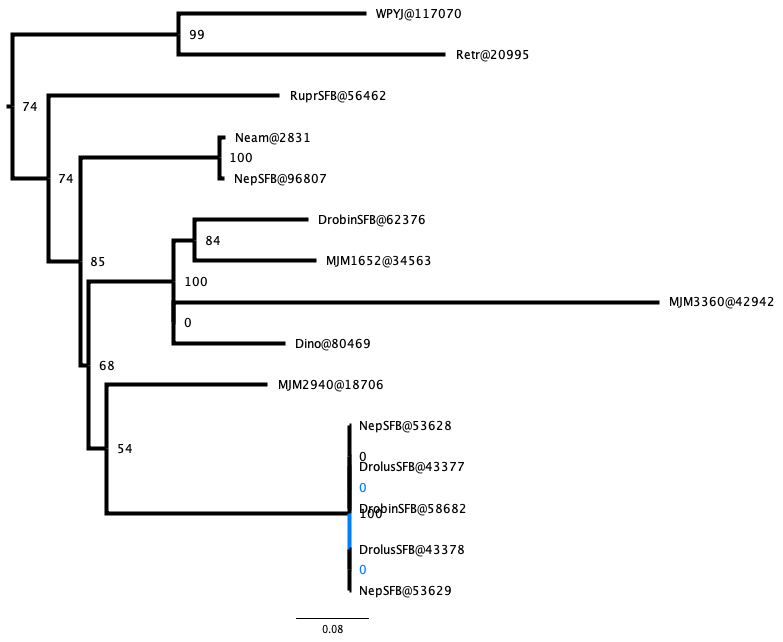

### Xyloglucan endotransglucosylases/hydrolase (cluster 1112)

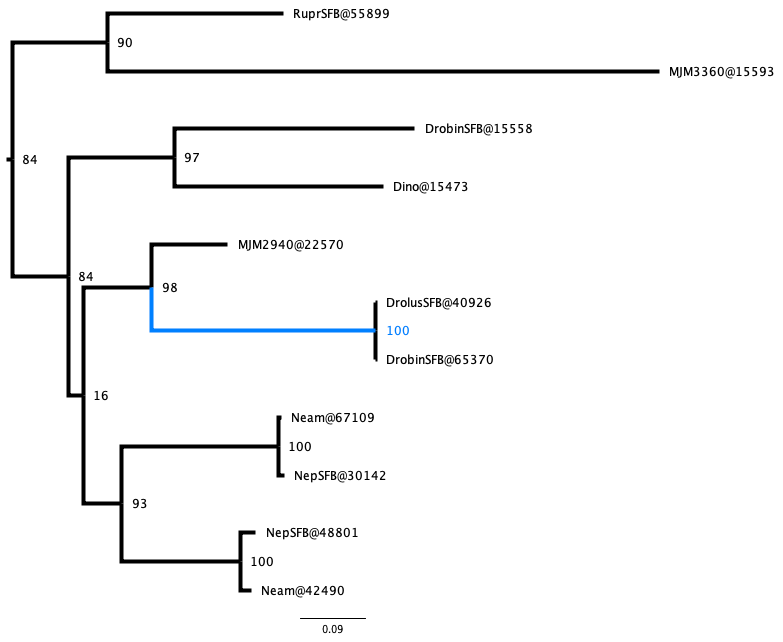

### Xyloglucan endotransglucosylases/hydrolase (cluster 3821)

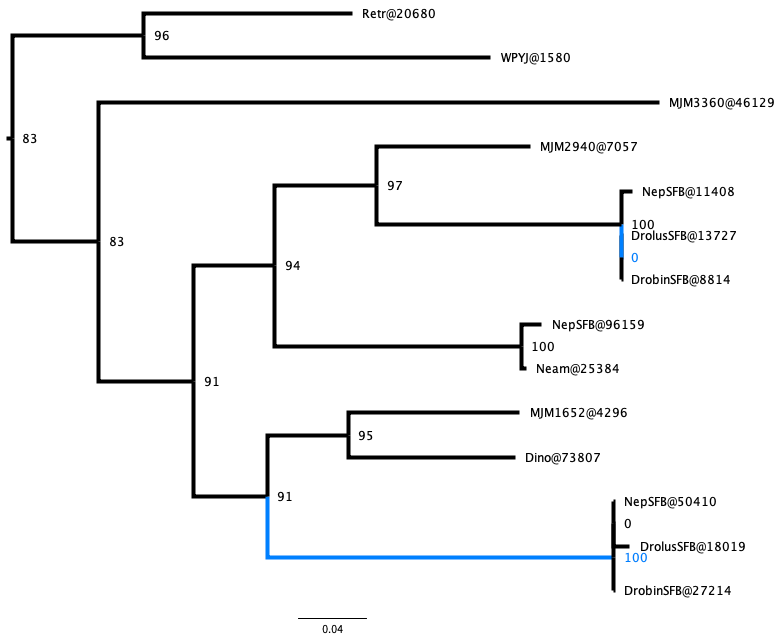

### Chitinase (cluster 2518)

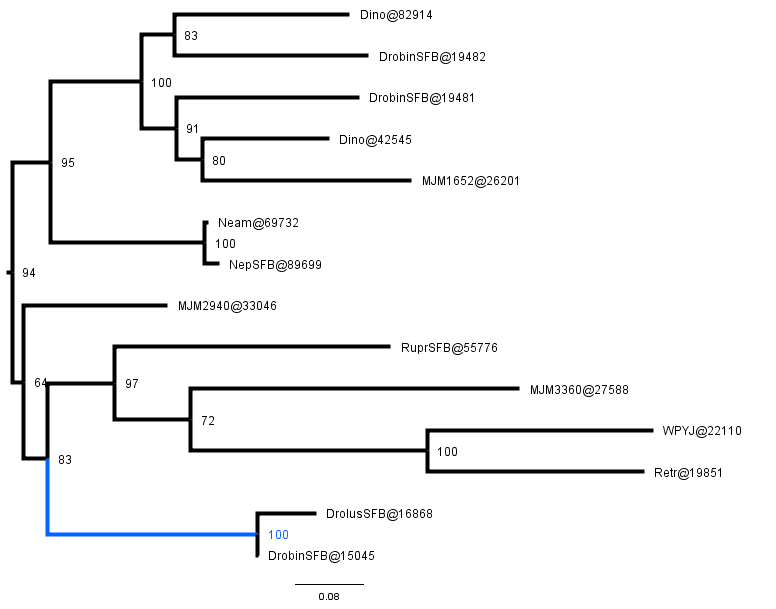

### Formate dehydrogenase (cluster 2136)

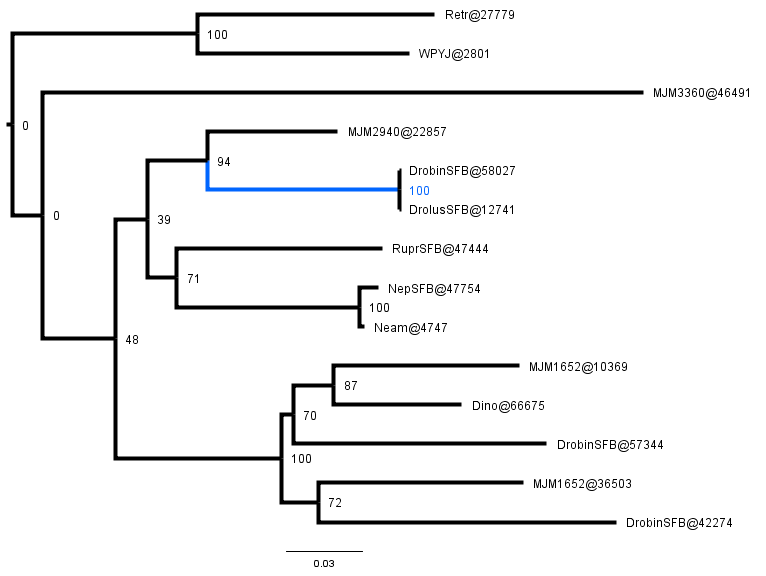

### Fructose-biphosphate aldolase (cluster 160)

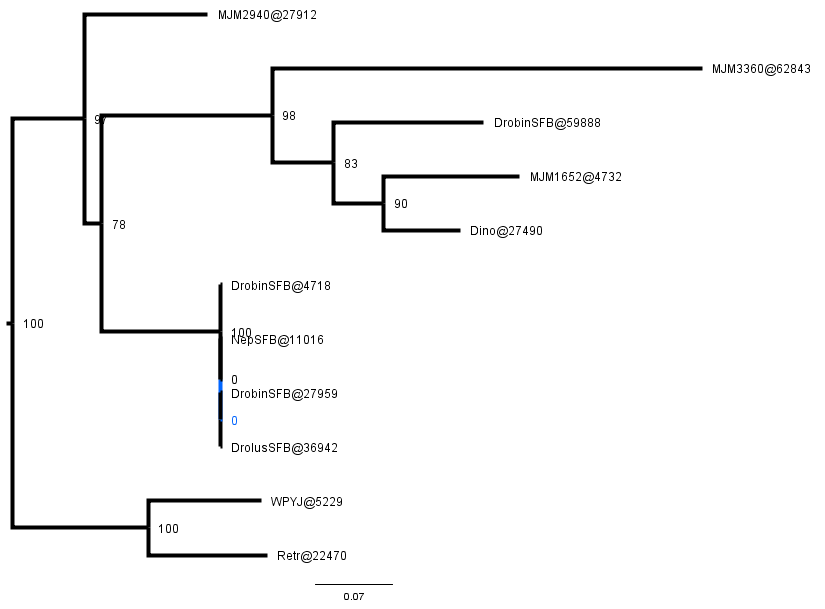

### Fructose-biphosphate aldolase (cluster 4896)

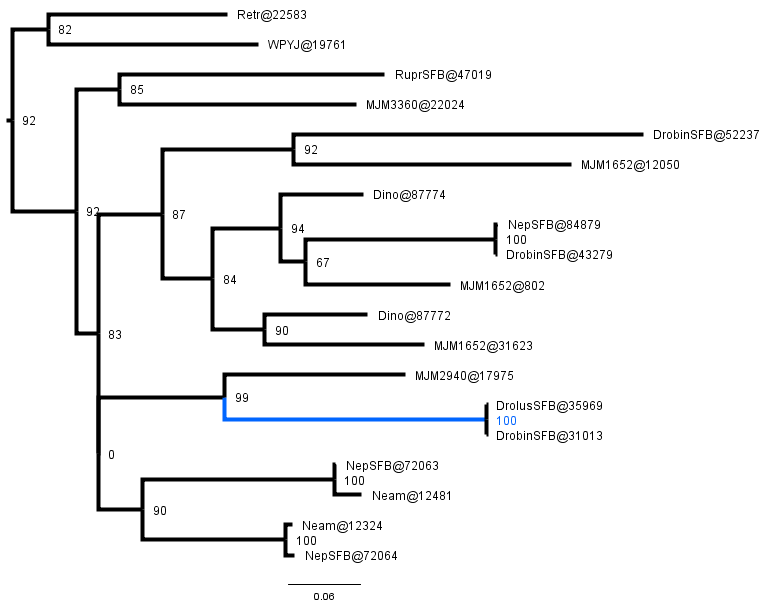

### Peroxidase (cluster 435)

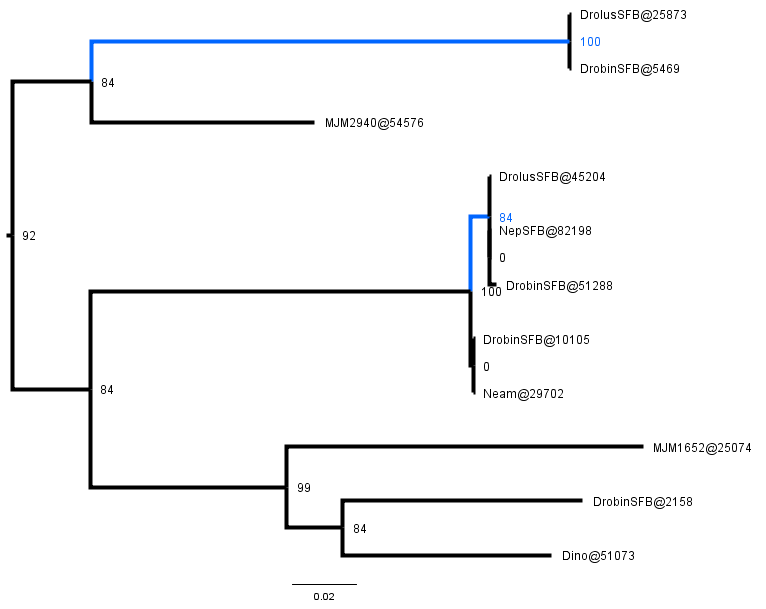

### Peroxidase (cluster 677)

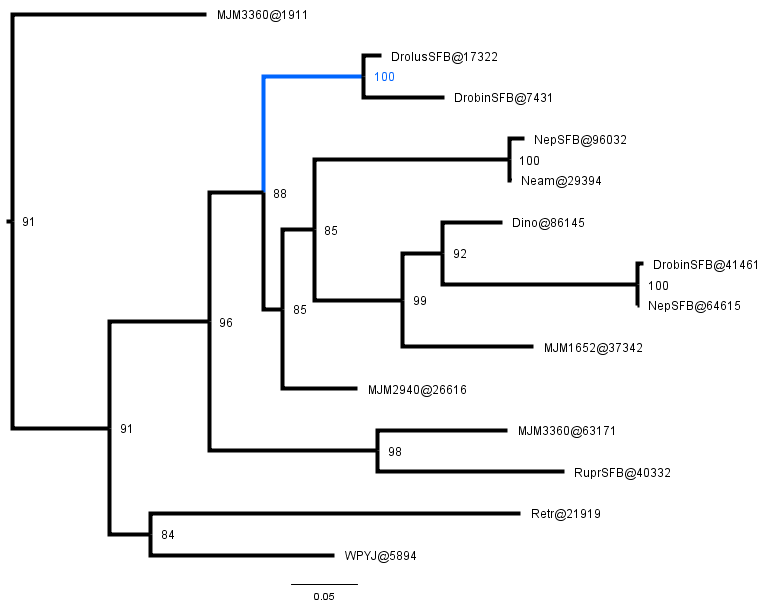

### Peroxidase (cluster 1196)

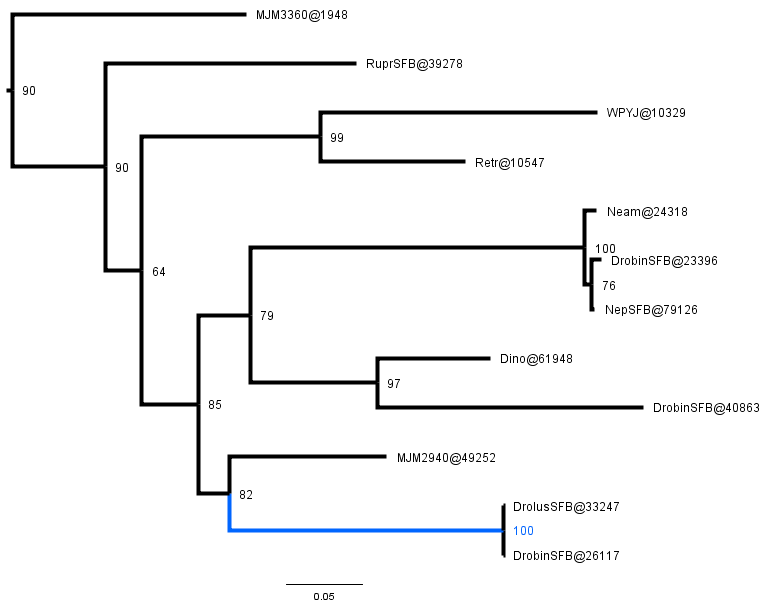

### Superoxide dismutase (cluster 3495)

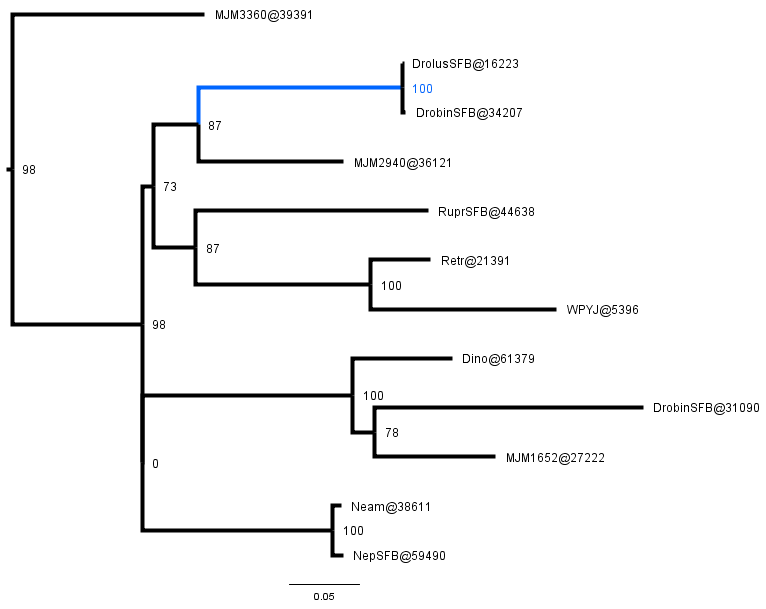

### Superoxide dismutase (cluster 3714)

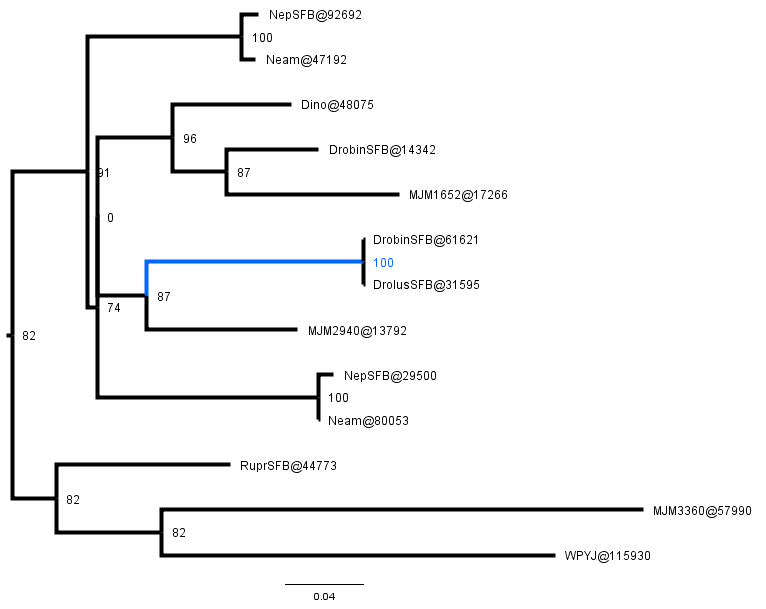

### Superoxide dismutase (cluster 999)

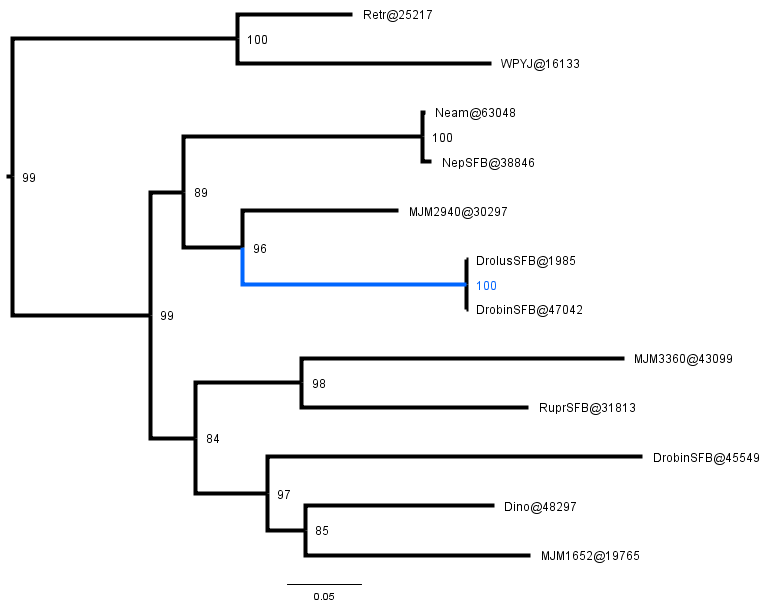

### Dicarboxylate transporter 1 (cluster 4294)

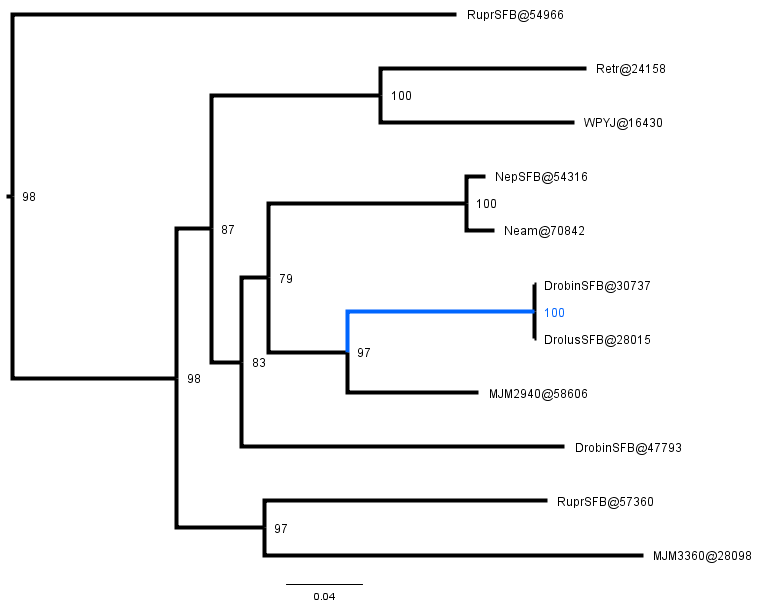

### Glutamine synthetase (cluster 926)

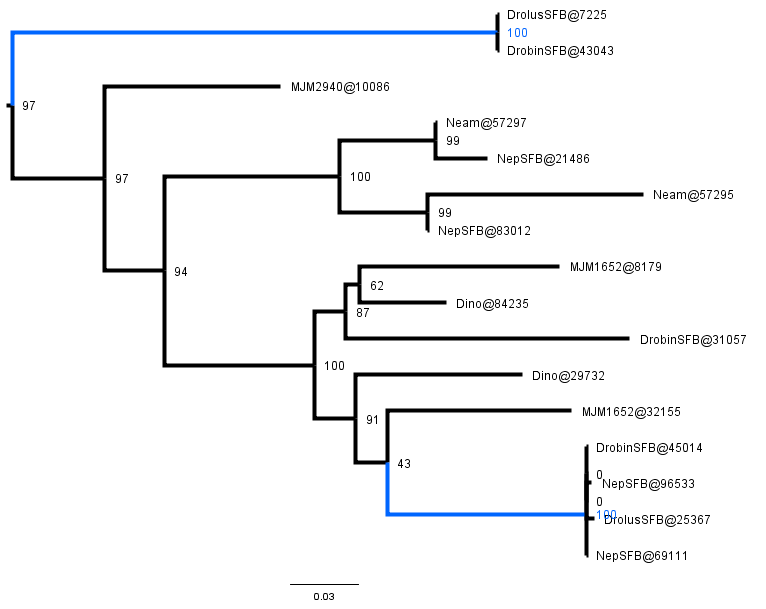

### Glutamine synthetase (cluster 315)

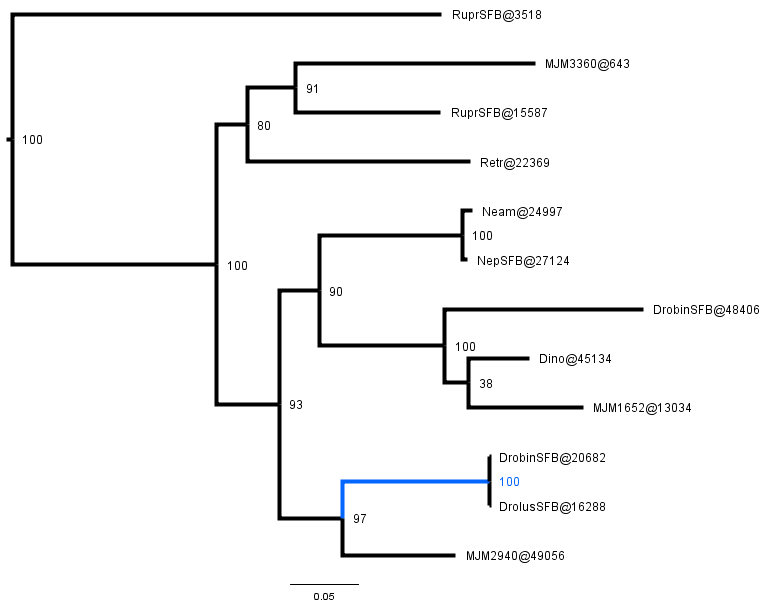

### Malate dehydrogenase (cluster 478)

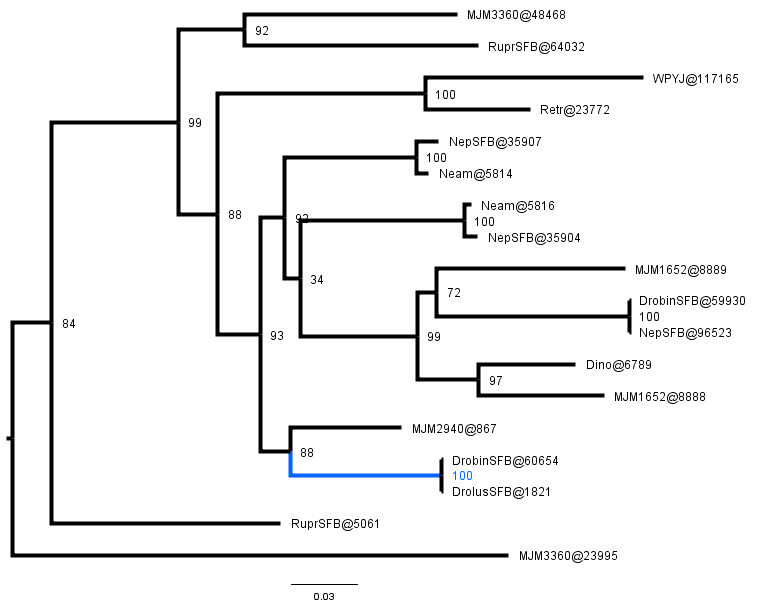

### Malate dehydrogenase (cluster 632)

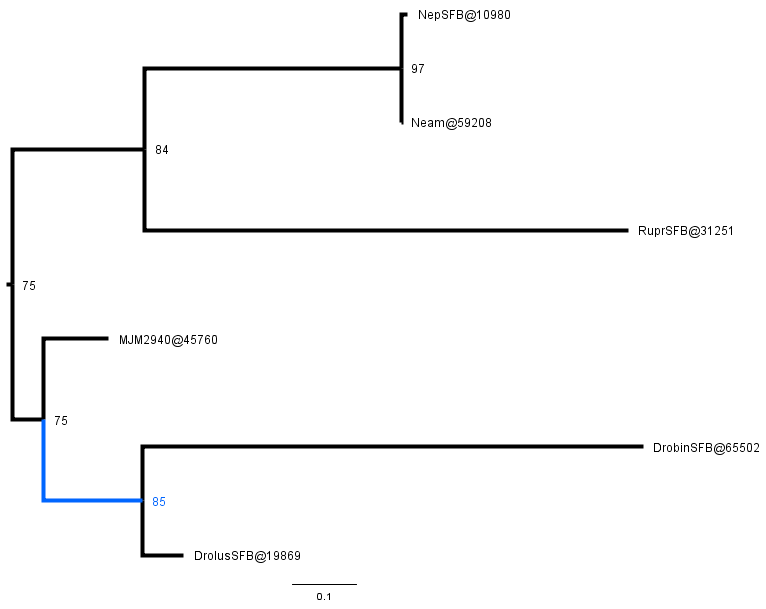

### *LBD4*-like (cluster 8006)

**Figure S4 Example of examining the relationship between a homolog tree and a species tree.** Nodes on the homolog gene tree where downstream taxa between the sister clades overlap are labelled as duplications. All non-duplication nodes are divided into their representative ortholog bipartitions and analysed for concordance or conflict with the species tree. The results of the conflict analysis between the homologous tree and the species tree are shown on the left. Concordance (blue) and conflict (red) are denoted at each node (Concordant/Conflict), and species tree bipartitions are shown on the right of the species tree.

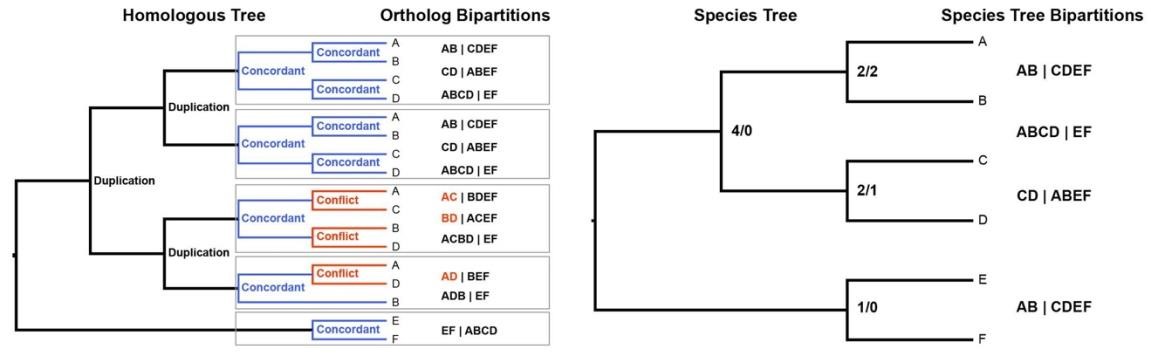

**Fig. S5 Representative picture of young leaf and mature trap tissues used for qRT-PCR assay.** (**A**) *Drosera coccicaulis* plant used to sample tissue for gene expression analyses presented in Fig. 5. RNAs were extracted from (**B**) unfurled leaf primordia (less than a 1cm long, trichomes absent or not yet able to secrete mucilage) and (**C**) mature leaves displaying fully developed sticky traps, densely covered with trichomes (red stalked glandular hair) secreting mucilage droplets. Scale bars = 5mm (A), 1mm (B), 5 mm (C).

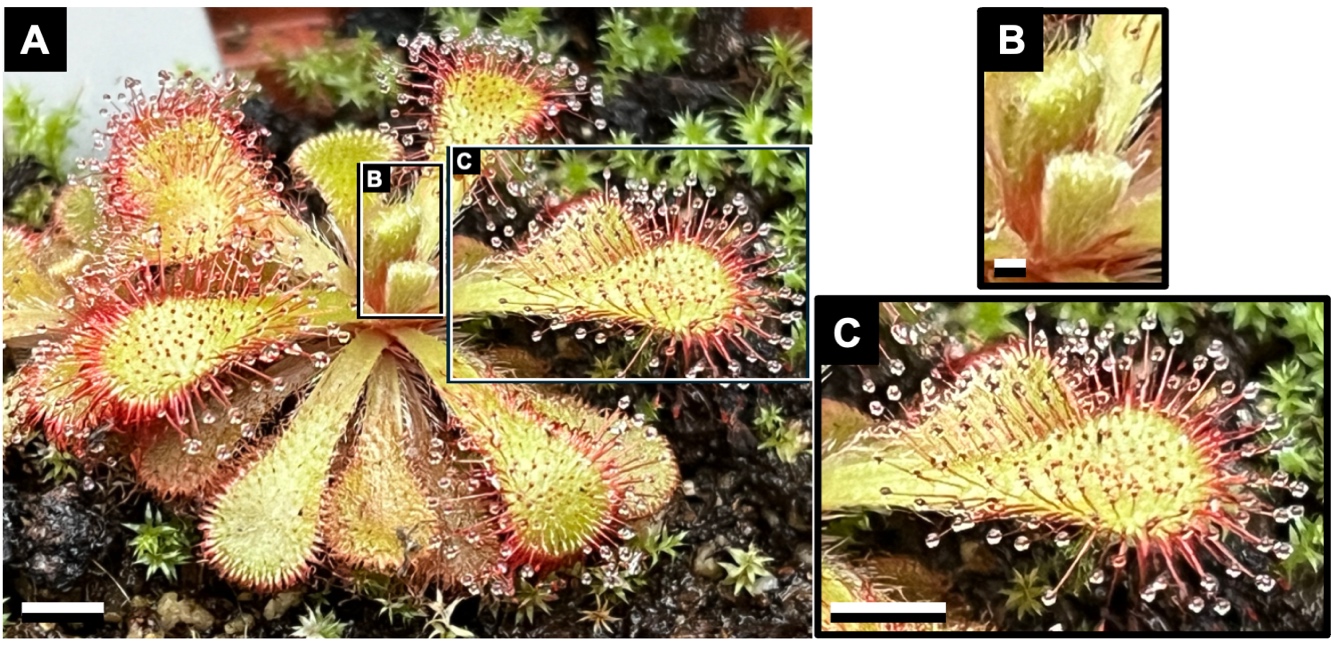

**Supplementary Method – Ancestral state reconstruction.**

We collected gene clusters for the families Ancistrocladaceae, Dioncophyllaceae, Droseraceae, Drosophyllaceae, Frankeniaceae, Nepenthaceae, Plumbaginaceae, Polygonaceae, and Tamaricaceae from GenBank on 16/09/2022 using PyPHLAWD (Smith and Walker 2019). The genes *matK* and *rbcL*, and the 18SrRNA were chosen to construct the tree to maximise species coverage, and additional sequences were manually downloaded from GenBank. Clusters were aligned and merged using MAFFT v7.471 (Katoh and Standley 2013). The sequence with the greatest number of characters was retained for each gene while lower-quality duplicates were discarded using a script written in Python [(https://github.com/HollyMaeRobertson/treebuildingscripts/blob/main/remove_repeats.py)](https://github.com/HollyMaeRobertson/treebuilding-scripts/blob/main/remove_repeats.py). The genes were concatenated using the program pxcat (v.0.9) from the Phyx package (Brown et al. 2017). We used RAxML-NG v. 1.1.0 (Kozlov et al. 2019) to build a tree from the concatenated genes, using a constraint tree of the known relationships of the non-core Caryophyllales (Walker et al. 2018), under the GTR+F+R5 model, which was chosen using the ModelFinder utility implemented in IQTREE v. 1.6.12 (Nguyen et al. 2015; Kalyaanamoorthy et al. 2017). Parsimony-based ancestral state reconstruction was performed on the carnivorous clade of the tree in Mesquite version 3.70 (Maddison and Maddison 2021).

**Supporting References**

Brown J.W., Walker J.F., Smith S.A. 2017. Phyx: phylogenetic tools for unix. Bioinformatics. 33:1886–1888.

Kalyaanamoorthy S., Minh B.Q., Wong T.K.F., von Haeseler A., Jermiin L.S. 2017. ModelFinder: fast model selection for accurate phylogenetic estimates. Nat. Methods. 14:587–589.

Katoh K., Standley D.M. 2013. MAFFT Multiple Sequence Alignment Software Version 7: Improvements in Performance and Usability. Mol. Biol. Evol. 30:772–780.

Kozlov A.M., Darriba D., Flouri T., Morel B., Stamatakis A. 2019. RAxML-NG: a fast, scalable and user-friendly tool for maximum likelihood phylogenetic inference. Bioinformatics. 35:4453–4455.

Maddison W.P., Maddison D.R. 2021. Mesquite: a modular system for evolutionary analysis.

Nguyen L.-T., Schmidt H.A., von Haeseler A., Minh B.Q. 2015. IQ-TREE: A Fast and Effective Stochastic Algorithm for Estimating Maximum-Likelihood Phylogenies. Mol. Biol. Evol. 32:268–274.

Smith S.A., Walker J.F. 2019. PyPHLAWD: A python tool for phylogenetic dataset construction. Methods Ecol. Evol. 10:104–108.

Walker J.F., Yang Y., Feng T., Timoneda A., Mikenas J., Hutchison V., Edwards C., Wang N., 974 Ahluwalia S., Olivieri J., Walker-Hale N., Majure L.C., Puente R., Kadereit G., Lauterbach M., Eggli U., Flores-Olvera H., Ochoterena H., Brockington S.F., Moore M.J., Smith S.A. 2018. From cacti to carnivores: Improved phylotranscriptomic sampling and hierarchical homology inference provide further insight into the evolution of Caryophyllales. Am. J. Bot. 105:446–462.
